# Discovery of thyrocyte heterogeneity reveals an essential role of Notch signaling in thyroid function and mammalian homeostasis

**DOI:** 10.1101/2022.09.02.506441

**Authors:** Lluc Mosteiro, Thi Thu Thao Nguyen, Simona Hankeova, Mike Reichelt, Shannon M. Vandriel, Zijuan Lai, Feroza K. Choudhury, Dewakar Sangaraju, Binita M. Kamath, Alexis Scherl, Robert Piskol, Christian W. Siebel

## Abstract

The thyroid functions at the apex of a web of endocrine organs that control cell growth, differentiation and metabolic homeostasis. Thyroid dysregulation significantly impacts human health in myriad ways with thyroid diseases standing as the most common endocrine disorder. Despite the essential role of the thyroid in human health, a high-resolution view of the cellular composition as well as molecular mechanisms that govern function of this crucial organ have been lacking. Employing the first single-cell analyses of adult mouse thyroid, we here report the discovery of unexpected thyrocyte heterogeneity, specifically three distinct thyrocyte subtypes marked by different metabolic and Notch signaling patterns. Using a battery of pharmacologic and genetic methods, we find that selective inhibition of Notch ligands and receptors disrupts thyrocyte mitochondrial activity and ROS production, thus decreasing levels of circulating thyroid hormones, inducing hypothyroidism and disrupting whole-body thermoregulation. We find an enriched frequency of hypothyroidism in children with Alagille Syndrome, a genetic disorder marked by Notch loss-of-function mutations, suggesting that our Notch-thyroid mechanisms are relevant in humans and directly account for Alagille hypothyroidism. Overall, our work reveals that Notch, although classically described as a developmental pathway that determines cell fate, controls homeostasis and thermoregulation in the adult through a mitochondria-based mechanism in a subset of thyrocytes. Our fine-grained picture of the thyroid unveils a novel understanding of this key metabolic organ and provides clinically impactful insights into its pathological dysfunctions.

## INTRODUCTION

The thyroid produces thyroid hormones that regulate cell growth, differentiation, development and metabolic homeostasis in all tissues (Brent, 2012), including orchestrating a cross-tissue network to control thermoregulation (Iwen et al., 2018). Given such essential roles in whole-body physiology, thyroid dysfunction and hormone imbalance significantly impact human health (Brent, 2012), with thyroid diseases standing as the most common endocrine disorder (Biondi and Wartofsky, 2012). The two primary thyroid cell types are thyrocytes (follicular cells) that synthesize, store and secrete the thyroid hormones thyroxine (T4) and triiodothyronine (T3) and parafollicular cells that secrete calcitonin. Thyrocytes have been considered a slow-turnover homogeneous population of cells that synthesize thyroid hormones in a multi-step process involving the uptake of iodine through the symporter Slc5a5, luminal coupling of the iodine to thyroglobulin (Tg) by thyroid peroxidase (Tpo) and lysosomal cleavage of iodinated Tg to form T4 and T3.

Despite the pivotal role of the thyroid in controlling metabolism, the lack of detailed understanding of the signaling pathways and cellular identities that regulate its function have hampered advances in novel therapeutic strategies. For example, while the Mouse Cell Atlas (Han et al., 2018) and Tabula Muris (Schaum et al., 2018) provide compendiums of single cell transcriptomic data from more that 40 murine tissues and reveal complex tissue heterogeneity with a new level of specificity, they unfortunately neglect the thyroid. Here, we report unexpected thyrocyte heterogeneity, including a subpopulation with high Notch and mitochondrial activities that resemble ‘classical’ thyrocytes as well as two other relatively quiescent subpopulations with lower Notch and metabolic activities. Using a unique battery of therapeutic antibodies that selectively inhibit each of the distinct Notch receptors and ligands (Lafkas et al., 2015; Tran et al., 2013; Wu et al., 2010; Yu et al., 2020), we discover that Notch signals are required for thyrocyte mitochondrial activity and function. Perturbing Notch signaling profoundly impacts thyrocyte function, and through doing so, disrupts adult homeostasis by triggering a cascade of thermoregulation defects that appear relevant not only in mouse models but also in humans.

## RESULTS

### The thyroid is comprised of a heterogeneous population of thyrocytes with distinct metabolic and Notch activity profiles

We appreciated that the thyroid, despite its central importance in regulating metabolism and maintaining homeostasis, has remained poorly characterized, especially using modern methods that illuminate fine resolution views of the cellular populations comprising a tissue. Thus, we performed the first single-cell transcriptomic analyses of murine thyroids. Tracking the expression of previously curated markers, we identified 16 cell types that included endothelial cells, erythrocytes, T cells, myeloid cells, fibroblasts, smooth muscle cells, parathyroid cells and Schwann cells (Figures 1A, 1B and S1A). A set of 16 known thyrocyte markers revealed two predominant populations of thyroid follicular cells or thyrocytes (TFC1 and TFC3) and one minor population (TFC2), uncovering unexpected heterogeneity (Figures 1A, 1B and S1A). TFC1 cells expressed high levels of thyrocyte markers, whereas the majority of TFC2 and TFC3 cells expressed low or undetectable levels of many thyrocyte-specific genes, including *Tshr, Slc5a5, Duox2*, and thyrocyte transcription factors *Pax8, Nkx2-1, Hhex* and *Foxe1* (Figures 1C and S1B). Spatial transcriptomic analysis (Ståhl et al., 2016) of thyroid sections also pointed to two distinct thyrocyte populations that, although expressing similar levels of Tg and Tpo, differentially expressed thyrocyte markers (*Pax8, Nkx2-1, Tshr* and *Slc5a5*) (Figures 1D and S1C). Immunostainings in thyroid sections confirmed this heterogeneity, with thyrocytes showing different levels of Pax8 (Figure S1D and S1E) and Nkx2-1 (Figure S1E). We note that our spatial transcriptomics and stainings suggest a possible spatial enrichment of each main thyrocyte type in the left vs. right lobes (Figure 1D, S1C and S1D). However, validating this hypothesis and determining whether these differences relate to previous observations of thyroid lobe asymmetry (Albi et al., 2012) requires further experimentation.

**Figure 1.**
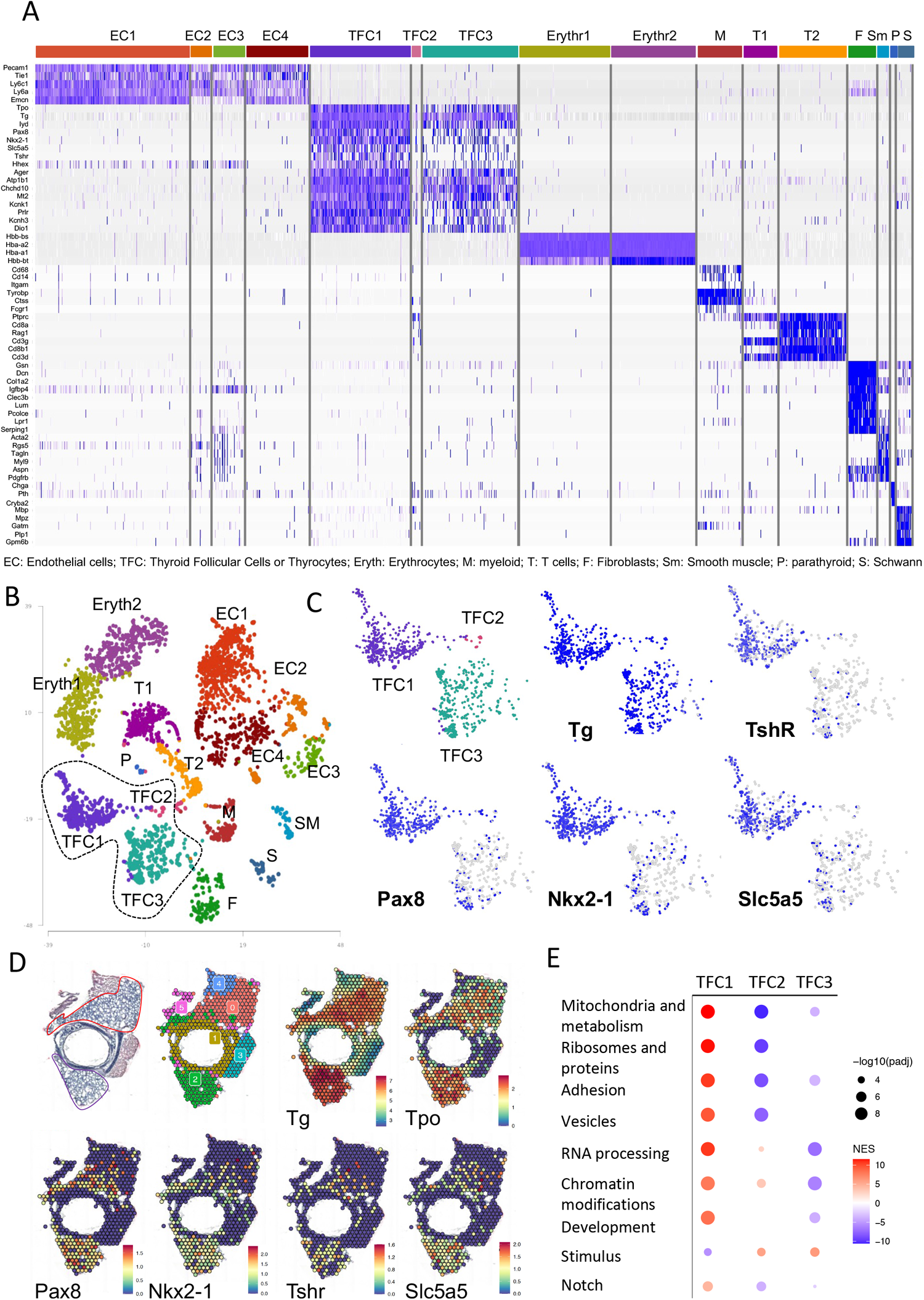
The thyroid is composed of a heterogeneous population of thyrocyte populations with distinct Notch activity profiles. **(A)** Heatmap of the z-scores of the expression of cell-specific markers used for cluster identification. The cluster identity is shown on the X axis and the gene names on the Y axis. **(B)** t-distributed stochastic neighbor embedding (tSNE) plot for all sequenced cells isolated from the thyroid of mice treated for 7 days with isotype-control antibody (n = 3, 40 mg/kg), aJ12 (n = 3, 20 mg/kg each) or aN12 (n = 3, 5 and 10 mg/kg, respectively). Colors highlight the 16 clusters identified: endothelial cells, EC; Thyroid Follicular Cells or thyrocytes, TFC; erythrocytes, Eryth; myeloid cells, M; T cells, T; fibroblasts, F; smooth muscle cells, SM; parathyroid cells, P; Schwann cells, S. **(C)** t-distributed stochastic neighbor embedding (tSNE) plot for thyrocyte clusters from (A) (TFC1, TFC2 and TFC3), with blue marking the cells expressing the indicated markers. **(D)** Visium spatial transcriptomic analysis showing haematoxylin-eosin staining, the Seurat integrated clusters (resolution 0.3) and the average expression per voxel of the indicated markers in thyroid sections of control mice, n = 2. Heatmap legends indicate normalized count per voxel. **(E)** Dotplot of the Gene Ontology (GO) terms and pathways enriched in each thyrocyte cluster. Colors represent the direction according to the normalized enrichment score (NES) (upregulated, red; downregulated, blue) and the dot sizes represents the −log10 (adjusted p-value, padj.). See also Figure S1.

Differential expression analyses defined gene sets that identified each thyrocyte cluster: 162 genes were upregulated in TFC1 compared to 122 genes downregulated in TFC2 and only 6 genes downregulated in TFC3 (Figure S1F). Pathway analysis indicated that the TFC1 gene set reflected upregulation of mitochondria-related pathways, whereas the TFC3 set indicated downregulation of RNA processing, chromatin modification and development pathways (Figure 1E). These data suggest that TFC1 comprises the metabolically active classical thyrocytes, whereas TFC2 and TFC3 seem comprised of underappreciated thyrocyte sub-types that are metabolically less active. We speculate that TFC3 may serve as a reservoir of progenitor-like cells, based on downregulation of pathways related to development and enrichment of several genes previously associated with stemness, such as *Tpt1* (Amson et al., 2013)(Figures 1E and S1G).

Markers of the developmental pathway Notch were also upregulated in TFC1 compared to TFC2 and TFC3 (Figure 1E). Notch signaling determines cell fate decisions during the development of many tissues, including the thyroid (Porazzi et al., 2012). Notch also controls the fate and function of adult cells in response to damage and under homeostasis (Siebel and Lendahl, 2017). In mammals, binding of cell surface ligands of the Jagged (Jag) or Delta-like (Dll) families to one of four Notch receptors (Notch1-4) (Gordon et al., 2015) triggers activation through a gamma-secretase-catalyzed proteolytic cleavage (Strooper et al., 1999). We analyzed the expression of Notch ligands and receptors in thyrocytes and observed that Jag1, Notch1 and Notch2 were enriched in TFC1 (Figure S1H), while only a few cells in TFC1 express Jag2, Notch3 and Dll ligands. The expression of a set of common Notch target genes (Stoeck et al., 2014) *Nrarp, Hes1, Hey1* and *Heyl* also seemed enriched in TFC1 cells (Figure S1H). RNA-scope *in situ* hybridization confirmed that some but not all thyrocytes express *Jag1, Notch1* and *Notch2* mRNA whereas *Jag2* mRNA expression appeared limited to a smaller subset of cells (Figure S1I), in agreement with our single cell data. Notch1 activity, assessed using an antibody that detects the gamma-secretase-cleaved (active) NICD1*, was restricted to a small number of cells that did not express the thyrocyte marker PAX8 nor the parafollicular cell marker calcitonin. Since these cells express endomucin (Figures S1J and S1K) and Notch1 commonly signals in endothelial cells (Kaspari et al., 2020), we conclude that the NICD1*-positive cells are endothelial cells. Notch2 activity, assessed through nuclear localization of NICD2, appeared clearly visible in thyrocytes (Figure S1K), with a heterogeneity of staining intensities from cell to cell, consistent with our single cell data and confirming that Notch2 is active in a subset of adult thyrocytes. Altogether, our single cell analyses reveal that the mouse thyroid contains three distinguishable thyrocyte populations, with the two main subtypes differing in expression levels of markers that define (a) classically-described thyrocytes, (b) thyrocyte function and metabolic activity and (c) the Notch signaling pathway.

### Notch is active in a subset of thyrocytes and its blockade induces thyrocyte defects

To better understand Notch signaling in the adult thyroid, we used monoclonal therapeutic antibodies, designed to selectivity inhibit individual receptors or ligands (Lafkas et al., 2015; Tran et al., 2013; Wu et al., 2010; Yu et al., 2020). Treatment with blocking antibodies against Notch receptors abolished Notch2 activity in thyrocytes and Notch1 activity in endothelial cells, demonstrating an effective signal blockade *in vivo* (and validating the activity detection methods described above) (Figures 2A, S2A). Treatment with the Jag1/2 blocking antibodies significantly reduced Notch2 signaling in thyrocytes (Figure 2A) but did not disrupt endothelial Notch1 activity (Figure S2A), consistent with the notion that Dll4 functions as the dominant Notch1 ligand in endothelial cells (Mack and Iruela-Arispe, 2018). Together with the ligand and receptor expression patterns described above, these results indicate that Jag1 and/or Jag2 induce Notch2 signaling in a subset of thyrocytes.

**Figure 2.**
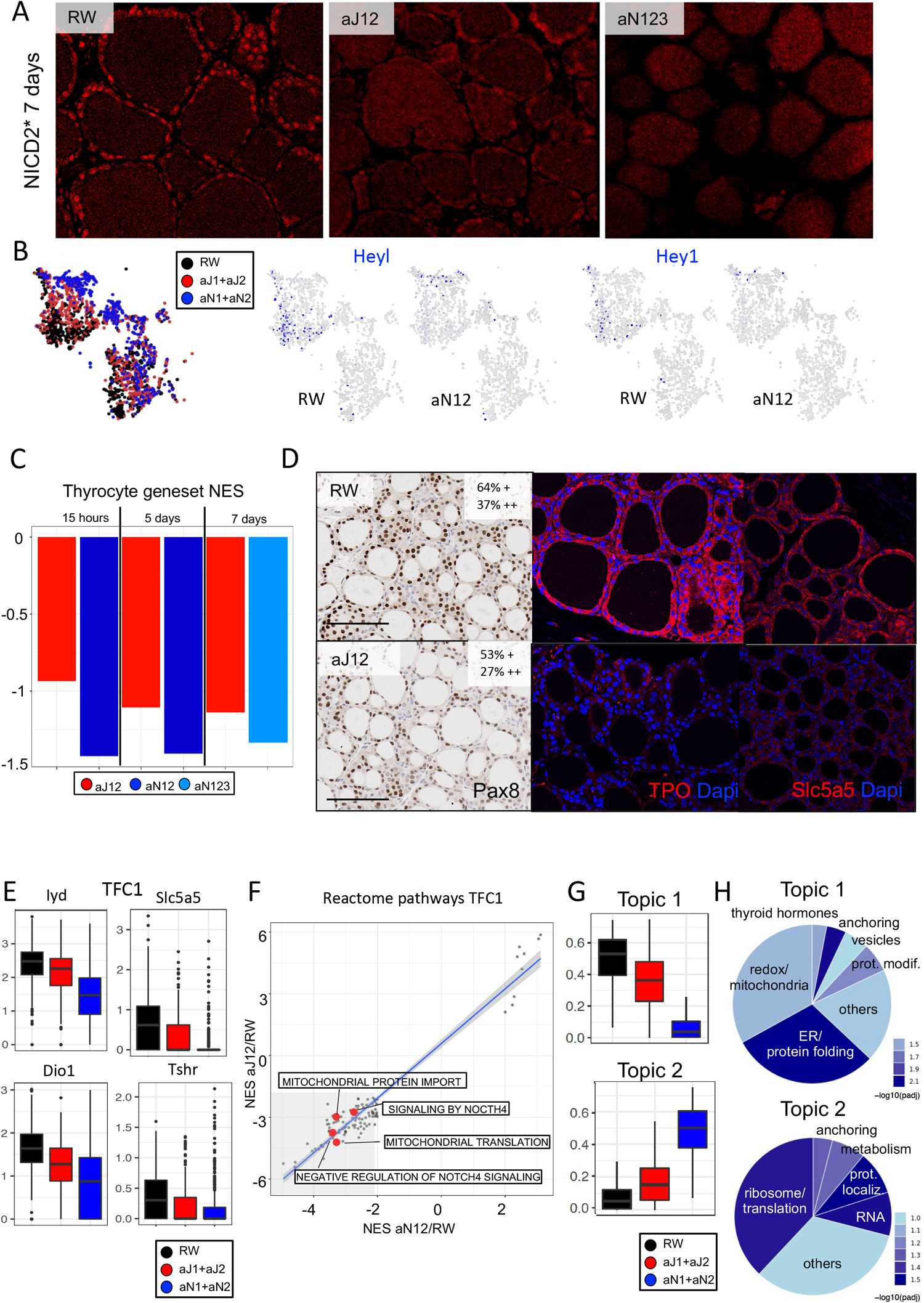
Notch activity in adult thyrocytes is essential for their function. **(A)** Representative images of immunofluorescent detection of gamma-secretase-cleaved (active) NICD2* in thyroid sections from mice (n = 5) treated for 7 days with isotype control aRW (40 mg/kg), aJ12 (20 mg/kg each), or aN123 (5, 10 and 20 mg/kg of each antibody, respectively). 40X objective. **(B)** tSNE plots for thyrocyte clusters (TFC1, TFC2 and TFC3) from mice treated as in (A) with the indicated antibodies showing cell clusters after each treatment (left panel) or cells expressing HeyL and Hey1 (right panels). **(C)** Normalized enrichment score (NES) of a thyrocyte gene set (Nkx2.1, Foxe1, Pax8, Hhex, Slc5a5, Iyd, Dio1, Tpo, Glis3, Cdca8, Slc26a4, Tshr) from thyroids (n = 5) isolated at the indicated times after treating mice as in (A) and (B). **(D)** Representative images of PAX8 immunohistochemistry and Tpo plus Slc5a5 immunofuorescence in thyroid sections from mice (n = 5) treated for 5 days as in (A). + +, strong positive; +, weak positive. Scale bars = 100mm; 40X objective. **(E)** Distribution of expression of the indicated genes in TFC1 cells from mice treated as in (B). n = 3 samples. **(F)** Gene Set Enrichment Analysis (GSEA) of Reactome pathways from cells in TFC1 from (B), highlighting in gray those with NES < −2 in both the aN12/aRW and aJ12/aRW comparisons. **(G)**Boxplots of Topics 1 and 2 enrichments in thyrocytes from mice treated as in (B). Y axis = topic scores. **(H)**Distribution of GO terms upregulated in Topics 1 and 2, colored by −log10(padj). Unpaired two-tailed Student’s t-test with Welch’s correction: *p*<0.05, *; *p*<0.01, **; *p*<0.001, ***; *p*<0.0001, ****. See also Figure S2.

To assess Notch function in thyrocytes, we repeated our single-cell analyses following Notch blockade. Notch blockade did not alter cluster composition, although cells from aRW, aJ12 and aN12 samples spatially segregated in both TFC1 and TFC3, suggesting transcriptional differences (Figure 2B). Notch blockade downregulated Notch target genes in TFC1, consistent with TFC1 as the Notch-active subpopulation and HeyL and Hey1 as downstream Notch targets (Figure 2B). RNA-scope also revealed that Notch blockade reduced thyroid expression of the Notch targets *Notch3, HeyL, Hey1* and *Nrarp* (Figures S2B and S2C). The anti-receptor antibodies induced greater effects than did the anti-Jag antibodies, reflecting our consistent experience of stronger Notch inhibition using anti-receptor versus anti-ligand treatments. Unbiased RNA-sequencing of bulk thyroid confirmed the significant downregulation of Notch target genes (Figure S2D) and pathway analysis pointed to Notch as one of the most downregulated (Figure S2E).

Strikingly, our sequencing analyses revealed that Notch blockade also reduced expression of the thyrocyte core transcription factors (*Pax8, Nkx2-1, Foxe1* and *Hhex*), as well as thyrocyte genes essential for the biosynthesis, storage and secretion of thyroid hormones, such as thyrotropin receptor *Tshr*, sodium/iodine cotransporter *Slc5a5*, iodo-tyrosine dehalogenase 1 (*Iyd*), dual oxidase *Dio1*, thyroid peroxidase (*Tpo*), *Glis3, Cdca8*, and pendrin (*Slc26a4*), while parafollicular genes were not affected (Figures 2C and S2F). We validated the downregulation of Notch target genes and thyrocyte genes using qRT-PCR (Figure S2G). Immunostaining of thyroid sections revealed that, although expression of the parafollicular cell markers calcitonin and calcitonin-gene related peptide (CGRP) remained unchanged (Figure S2H), Notch inhibition decreased expression of the thyrocyte lineage transcription factor PAX8 (Figure 2D and S2I) as well as the peroxidase Tpo and the symporter Slc5a5 (Figure 2D), both essential for thyroid hormone production. The downregulation of thyrocyte genes was exclusively observed in TFC1 (Figure 2E and S2J). Notch inhibition also decreased nuclear PAX8 expression in the thyrocyte cell line FRTL5 (Figure S2K), confirming that Notch inhibition directly affects thyrocytes. We also exploited organoids---multicellular structures with polarized cells that recapitulate organ development, physiology and histological architecture---as an additional *in vitro* model. Control organoids displayed organized structures resembling thyroid follicles that expressed markers of both thyrocytes and parafollicular cells, suggesting some degree of bipotency or plasticity (Figure S2L). In contrast, we observed that JAG blockade dramatically reduced the frequency of organoid formation (Figure S2M); furthermore, those organoids that grew expressed little PAX8 protein and were less structured and smaller (Figures S2M and S2N). Together, these data demonstrate that a subset of adult thyrocytes maintain Notch activity to sustain a gene expression program essential for thyrocyte function; inhibiting this program induces thyroid defects that manifest as defective organoid formation *in vitro* and decreased thyrocyte marker expression *in vivo*.

To investigate mechanisms through which Notch inhibition disrupts thyroid function, we analyzed the downstream transcription changes. In TFC1, Notch inhibition upregulated 578 genes and downregulated 496 genes, whereas few effects were observed in TFC2 cells (10 genes downregulated), which do not express detectable levels of Notch genes. Notch blockade in TFC3 cells induced upregulation and downregulation of 116 and 64 genes, respectively, although the low expression of Notch ligands and receptors in these cells suggests that such changes may be indirect and reflect Notch signal inhibition in another cell type(s). GSEA analysis showed that Notch blockade downregulated pathways related to mitochondria in TFC1 thyrocytes (Figure 2F) and posttranslational modification in TFC3 (Figure S2O). To better resolve the molecular differences between treatments, we used topic modeling, which infers transcriptional programs within cells and correlates them with transcription factor activities. After evaluating 2-15 topics for their fitness to the data, we chose seven topics to optimally represent the transcriptional programs that most impactfully distinguish prominent cell types (Figure S2P). By mapping topic scores onto thyrocyte tSNE plots (Figure S2Q), we discovered that Notch blockade inhibited the topic 1 program, defined by Redox/mitochondrial pathways, and stimulated the topic 2 program, related to ribosomes and translation (Figures 2G and 2H). These results agreed with the GSEA analysis in TFC1, which also showed that Notch blockade downregulated pathways related to mitochondria (Figure 2F) and further suggest that Notch regulates mitochondrial activity in TFC1 thyrocytes.

### Notch blockade induces mitochondrial defects in thyrocytes

Given our sequencing, topic and pathway analyses---together with the knowledge that thyroid hormone synthesis demands robust mitochondrial activity to generate reactive oxygen species (ROS) (Szanto et al., 2019)---we investigated whether mitochondrial dysfunction might explain the essential role of Notch in thyrocytes. Notch inhibition downregulated expression of multiple mitochondrial genes in the TFC1 cluster (in a manner that correlated with the magnitude of inhibition, anti-N1/2 > aJ1/2; Figure S3A). Transmission electron microscopy revealed striking structural defects in the mitochondria from thyrocytes but not in other thyroid cell types (Figures 3A and S3B), revealing selectivity in the cell type affected. Whereas control thyrocytes showed a typical condensed mitochondrial matrix with evenly-spaced and well-defined cristae, Jag1/2 blockade correlated with aberrant mitochondria that displayed a brighter matrix, irregular or missing cristae, and swelling at the mitochondrial ends, phenotypes that were exacerbated after Notch1/2 inhibition (Figure 3A). Notch inhibition also led to mitochondrial membrane depolarization in primary thyrocytes, based on loss of TMRM fluorescence in the polarized mitochondrial membrane (Gerencser et al., 2012), to a similar extent as did the membrane-depolarizing control compound THPB (Figures 3B and S3C). Likewise, Notch inhibition reduced activity of Complex IV (Cox4), which is required for membrane polarization (Reguera et al., 2020), based on Cox4 staining and *ex vivo* assays of Cox4-induced cytochrome C oxidation (Figures 3C, 3D, S3D and S3E). Mitochondria content, assessed by the ratio of mDNA/gDNA was not affected, indicating that these changes did not reflect a decrease in mitochondria number (Figure S3F). To better assess whether Notch directly impacted thyrocyte mitochondrial function, we tested the FRTL5 thyrocyte cell line using (a) the Cell Mito Stress assay to calculate the oxygen consumption rate after inhibiting key components of the mitochondrial electron transport chain and (b) the ATP Real-Time assay to calculate mitochondrial and glycolytic ATP production rates after metabolic modulation. Notch inhibition perturbed mitochondrial function based on decreases in multiple parameters, including basal mitochondrial respiration, maximal respiration, spare capacity and ATP-linked respiration and production, without altering non-mitochondrial CO_2_ consumption (Figures 3E and S3G).

**Figure 3.**
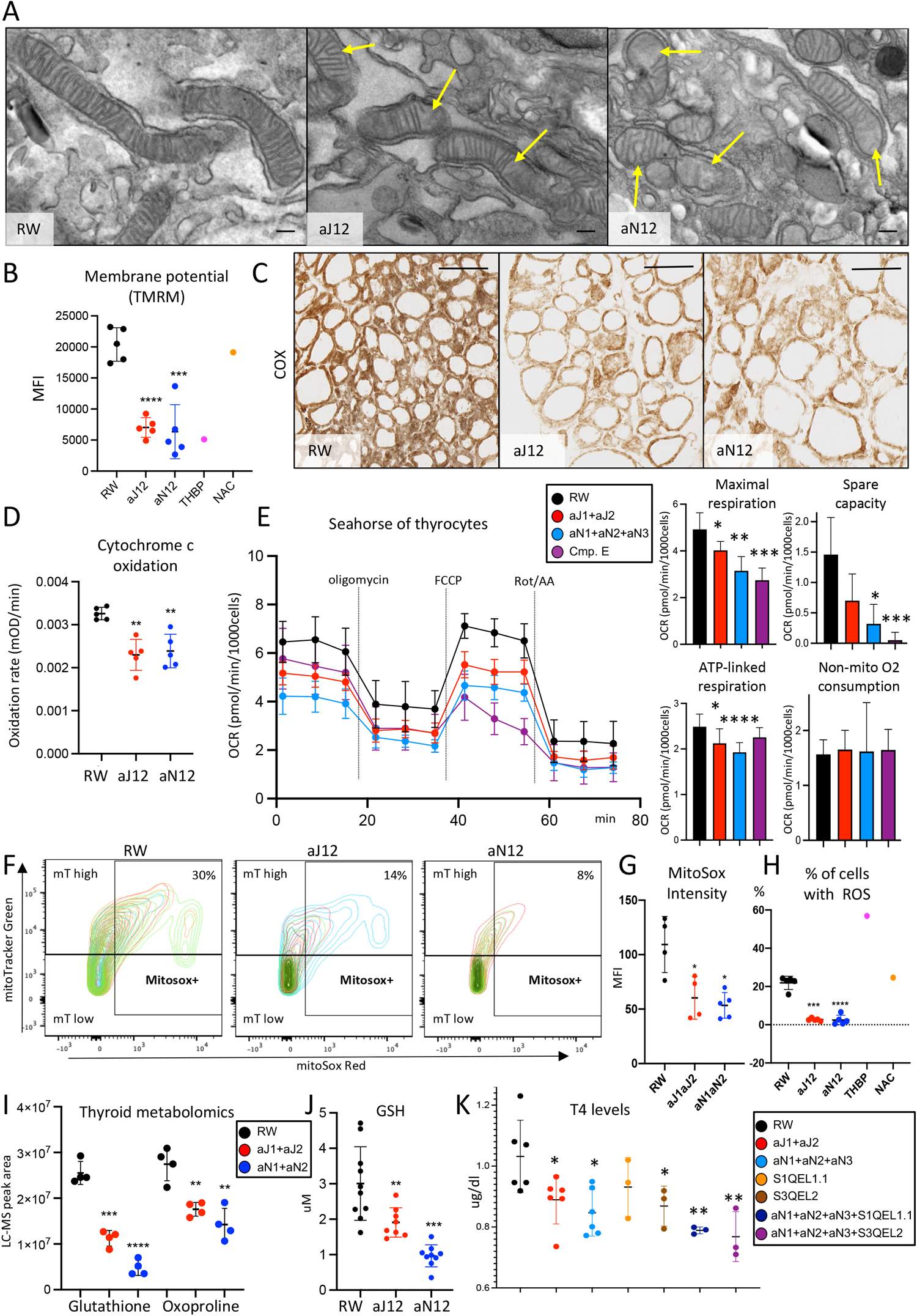
Notch blockade induces mitochondrial defects in thyrocytes. **(A)** Representative scanning electron microscopy images of thyrocyte mitochondria from mice (n = 3) treated for 7 days with aRW (40 mg/kg), aJ12 (20 mg/kg each) or aN12 (5 and 10 mg/kg of each antibody, respectively). Arrows = aberrant mitochondria. Scale bars = 200nm. **(B)** Mean fluorescence intensity of TMRM to measure mitochondrial membrane potential in primary thyrocytes isolated from mice (n = 5) treated as in (A). THBP (100uM) or NAC (10mM) treatments *in vitro* served as positive or negative controls, respectively. Ave. ± s.d. **(C)** Cox staining (VitroView™) in thyroid cryosections from mice (n = 4) treated as in (A) Scale bars = 100mm. **(D)** Cytochrome c oxidation rate in primary thyrocytes isolated from mice (n = 5) treated as in (A). Ave. +/-s.d. **(E)** Cell Mito Stress Seahorse assay in the FRTL5 thyrocyte cell line cultured for 2 days with aRW (25μg/ml), aJ12 (25μg/ml each), aN123 (25μg/ml each) or Compound E (Cmp. E) (1μM). Oligomycin inhibits ATP synthase (complex V), decreasing electron flow through the electron transport chain (ETC) and the oxygen consumption rate (OCR) linked to ATP production. FCCP collapses the proton gradient, allowing complex IV to reach its maximum. Spare capacity = maximal – basal respiration. Roterone/AA inhibit complexes I and III, respectively, shutting down mitochondrial respiration without affecting non-mitochondrial respiration. Ave. ± s.d.; n = 5 wells. **(F)** Fluorescent density intensity plots following 1h staining with mitotracker Green (mT), which accumulates in the mitochondria, and MitoSox red, which reacts with superoxide (O_2_^-^), from thyrocytes isolated from mice (n = 5) treated as in (A). **(G)** Mean fluorescence intensity of MitoSox red staining in (F). Ave. ± s.d. **(H)** Percentage of positive cells following 1h DCFDA staining of thyrocytes isolated from mice (n = 5) treated as in (A). THBP (100uM) and NAC (10mM) served as controls as in (B). Ave. ± s.d. **(I)** Mass spectrometry measurements of the indicated metabolites in thyroids from mice (n=4) treated as in (A). Ave. ± s.d. **(J)** Glutathione (GSH) concentrations in thyrocytes isolated from mice (n = 8) treated as in (A) and measured by fluorometry Ave. ± s.d. **(K)** ELISA measurements of T4 concentrations FRTL5 thyrocyte cell culture medium following treatment as in (E) and including 1μM S1QEL1.1 or S3QEL2 as indicated. Ave. ± s.d.; n = 3-6 wells. Statistical significance was assessed using the unpaired two-tailed Student’s t-test with Welch’s correction: *p*<0.05, *; *p*<0.01, **; *p*<0.001, ***; *p*<0.0001, ****. See also Figure S3.

To investigate ROS production in the thyroid, we combined mitotracker Green (mT), which accumulates in the mitochondria, with mitosox Red, which reacts with superoxide (O_2_^-^). Notch inhibition reduced the percentage of thyroid cells with high mitotracker staining while concomitantly increasing the percentage with low mitotracker, confirming that Notch blockade decreases mitochondrial function (Figures 3F and S3H). Moreover, Notch blockade reduced superoxide generation as demonstrated by a decrease in mitosox intensity and double positive (DP) cells (Figures 3F, 3G and S3I). DCFDA staining for all reactive oxygen species (Maharjan et al., 2014) revealed decreased intensity and percentage of ROS+ cells (Figures 3H and S3J), demonstrating that the mitochondrial dysfunction induced by Notch inhibition results in decreased levels of all ROS, including SO3-.

Given that ROS changes could reflect Notch-controlled mitochondrial function or indirectly result from general changes in glycolysis that would limit essential mitochondrial metabolites, we performed thyroid metabolomics. Principal component analysis (PCA) of metabolomic data from isolated thyroids clustered samples according to anti-Notch treatments (Figure S3K). Notch inhibition had mixed effects on intermediates and metabolites related to glycolysis, glutamine synthesis and the urea cycle (Figures S3L-N). Ketogenesis metabolites increased, indicating that acetyl-coA was not limiting (Figure S3M). In contrast, we observed clear changes in oxidative stress, with reduced levels of metabolites such as glutathione and oxoproline (Figure 3I). Likewise, although catalase activity remained unchanged (Figure S3O), we observed decreases in GSH and GSSG (Figures 3J and S3P), consistent with decreased ROS generation. Compounds that suppress mitochondria complexes I (S1QEL1.1) or III (S3QEL2), which are required for superoxide production, led to reduced secretion of the thyroid hormone thyroxine or T4 in FRTL5 thyrocyte cells, confirming that mitochondrial activity is essential for T4 secretion (Figure 3K). Notch blockade caused a similar decrease in T4 secretion (Figure 3K and S3Q). These data thus support a model in which Notch inhibition directly disrupts thyrocyte mitochondrial function, resulting in decreased ROS production and T4 secretion.

### Notch inhibition triggers hypothyroidism and a cascade of thermoregulation defects

Having observed that Notch inhibition disrupted mitochondrial function and T4 secretion *in vitro*, we investigated whether Notch blockade also affected thyroid function *in vivo*. Histological examination showed that Notch inhibition reduced the number of re-uptake vacuoles and induced flattening of the cuboidal epithelium (Figures 4A and S4A), suggestive of decreased thyrocyte activity according to these standardized criteria (Lee et al., 2016). Indeed, we discovered notable decreases in the serum levels of T3 and T4 with both pan-Jag and pan-Notch inhibition (Figures 4B and S4B), with T4 levels dropping as early as 24 hours after anti-J1/2 treatment and progressively decreasing over seven days (Figure 4C). Inhibiting any single Notch ligand or receptor was insufficient to affect T4 levels; in contrast, inhibiting both Jag1 plus Jag2 or Notch1 plus Notch2 was necessary and sufficient to decrease T4 levels, with the combined receptor blockade showing the strongest defect (Figure S4B). Thyroid secretion of T4 is controlled by the hypothalamus and pituitary gland through the secretion of thyrotropin releasing hormone (TRH) and thyroid stimulating hormone (TSH). Lower T4 levels anti-correlated with elevated levels of TRH and TSH (Figure S4C and S4D), consistent with decreased thyroid secretion of T4 and T3 triggering a compensatory increase in TRH and TSH secretion (Sheehan, 2016). In contrast to T4 and T3, levels of calcitonin, which is secreted by parafollicular thyroid cells, remained unchanged after anti-J1/2 treatment (Figure S4E), confirming that thyrocytes are the relevant Notch-responsive thyroid cell type. Together, these data demonstrate that Notch activity is essential to maintain normal levels of circulating thyrocyte hormones. Notch blockade *in vivo* disrupts thyrocyte function and induces hypothyroidism.

**Figure 4.**
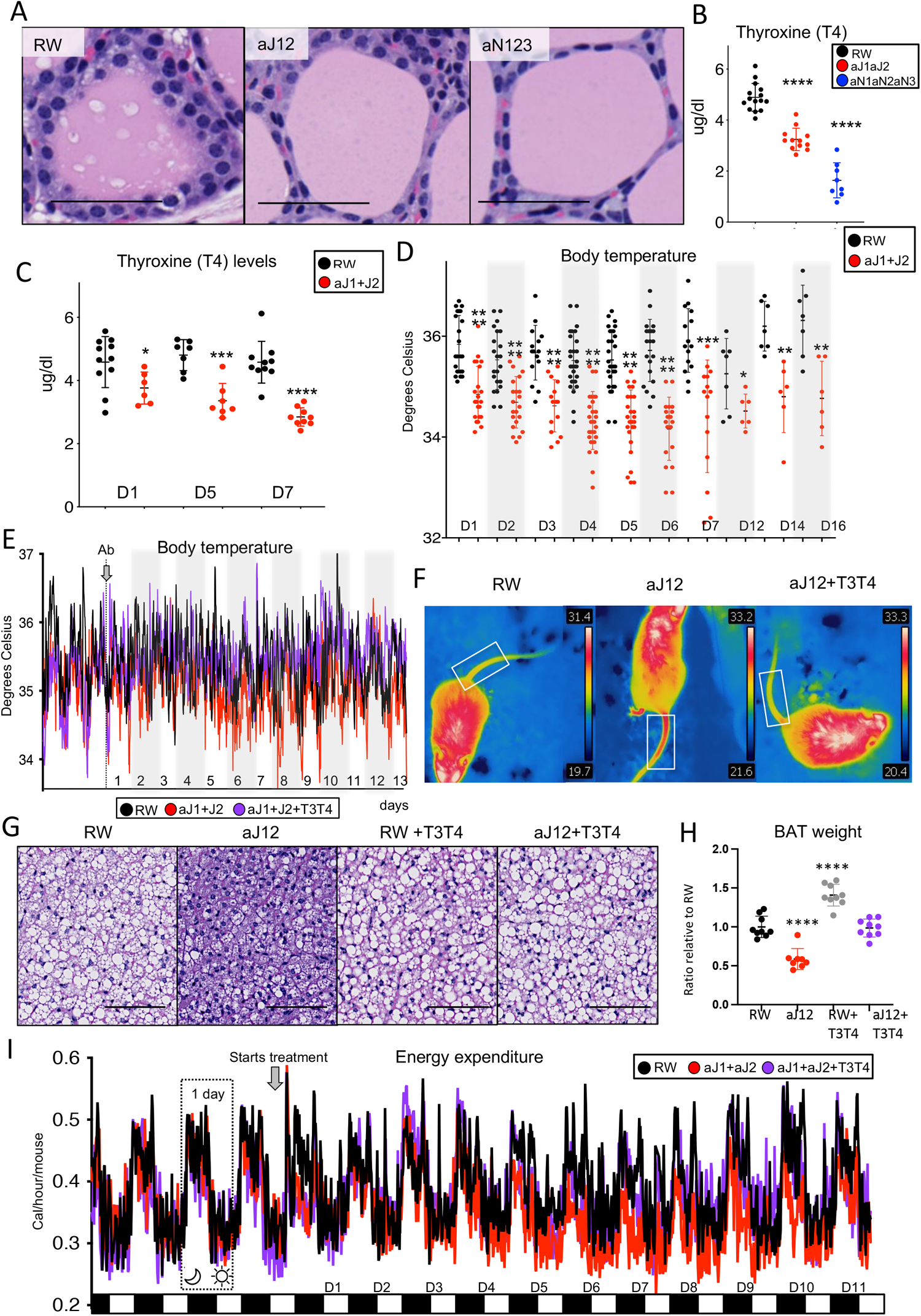
Notch-induced hypothyroidism triggers a cascade of thermoregulation defects. **(A)** Representative haematoxylin and eosin staining of thyroid sections from mice (n = 10) 7 days after a single i.p. injection of aRW (40 mg/kg), aJ12 (20 mg/kg each) or aN12 (5 and 10 mg/kg of each antibody, respectively). Scale bars = 100mm. **(B)** Thyroxine (T4) serum levels from mice (n ≥ 8) treated as in (A), measured by ELISA. 3 independent experiments, Ave. ± s.d. **(C)** T4 measurements as in (B) over the indicated time course (D = days). n ≥ 10 mice. **(D)** Body temperatures measured using a rectal probe at the same time on each of the indicated days in mice (n ≥ 6) housed at 22°C and treated as in (A). 5 independent experiments, Ave. ± s.d. **(E)** Body temperatures measured at 15 min intervals over the indicated days using telemetric transponders implanted in the abdominal cavities of mice (n = 5) treated as in (A), with the addition of a hormone rescue group of mice that were also dosed daily with T4 and T3 (1 and 0.05 μg/g, respectively). Ave. ± s.d. **(F)** Representative infrared thermographs of mice (n = 5) obtained after 4 days of treatment as in (E). Legend = maximum and minimum temperatures registered. **(G)** Representative haematoxylin and eosin staining of brown adipose tissue (BAT) sections obtained after 7 days from mice (n = 5) treated as in (E). Scale bars = 100mm. **(H)** Wet weight of brown adipose tissue (BAT) in mice (n = 9) treated as in (E), normalized to the average wet weight measured in the control group. Ave. ± s.d. **(I)** Energy expenditure in mice (n = 5) treated as in (E). Black and white bars represent daily night and day cycles, respectively. Oxygen and carbon dioxide levels were measured every 15 min for 3 days before and 11 days after treatment, as indicated. Ave. ± s.d. Statistical significance was assessed using the unpaired two-tailed Student’s t-test with Welch’s correction: p<0.05, *; p<0.01, **; p<0.001, ***; p<0.0001, ****. See also Figure S4.

Given the key role of the thyroid in regulating whole-body homeostasis, including thermoregulation and metabolism (Warner et al., 2013; Weiner et al., 2016), we evaluated whether Notch inhibition induced physiological defects consistent with hypothyroidism in mice (Ueta et al., 2011; Warner et al., 2013; Weiner et al., 2016). We focused on Jag blockade to avoid the intestinal and weight loss phenotypes of Notch1 plus Notch2 inhibition (Pellegrinet et al., 2011; Wu et al., 2010). Jag inhibition reduced body temperature by approximately 1°C as early as 24 hr after dosing, with temperature loss progressing during the first week, paralleling the decreases in T4 serum levels (Figures 4C and 4D). Previous research indicates that genetically obese mice (Ob/Ob) display decreased body temperature through mechanisms that at least partly stem from thyroid defects, including lower levels of thyroid hormones (Weiner et al., 2021). On the other hand, thermoneutrality (30°C) and moderate (22°C) or stringent (4°C) cold challenges can alter thermoregulatory responses, with hypothyroid animals succumbing rapidly after being moved to cold environments (Kaspari et al., 2020; Ring, 1942). We thus tested Notch blockade under these different genetic or temperature-challenge conditions, where we found that antibody treatment consistently decreased T4 (Figure S4F). Notably, exogenous administration of thyroid hormones (Figure S4G) restored body temperature to control levels at 22°C and under cold-challenge conditions, as assessed using telemetry and a rectal thermometer (Figures 4E, S4H and S4I). T3T4 administration did not affect mitochondrial morphology in aRW treated thyrocytes and did not rescue the aberrant mitochondrial morphology triggered by Notch inhibition (Figure S4J), suggesting that the mitochondrial defects are upstream of the thyroid hormone deficiency. These results together reveal that Notch blockade induces a drop in core body temperature that correlates with decreases in thyroid hormones and can be rescued by hormone supplementation.

Numerous physiological systems tightly control thermogenesis and core body temperature in warm blooded animals (Sanchez-Gurmaches et al., 2016). Using infrared thermography, we observed that Notch blockade increased heat dissipation from the tail, highlighting one mechanism consistent with a drop in core body temperature; furthermore, T3T4 rescued the heat dissipation phenotype, thus linking heat dissipation to thyroid hormone activity (Figures 4F and S4K), consistent with previous work (Warner et al., 2013). In response to a drop in core temperature, mammals also generate heat through BAT activation, previously linked to dysregulated thyroid function (Warner et al., 2013). Histological analysis following JAG blockade revealed numerous signs of BAT activation, including decreases in lipid droplets, adipocyte size and BAT weight, and T3T4 supplementation rescued these changes (Figures 4G, 4H, S4L and S4M). Energy dissipation in BAT is mediated by Ucp1, which was upregulated after aJ12 treatment at the mRNA (Figure S4N) and protein levels (Figure S4O). Other genes important for BAT thermogenic function, such as the *Ucp1* coactivator *Ppargc1*, as well as genes involved in BAT thermogenesis and fatty acid oxidation such as *Pparg, Cidea, Prdm16* or *Acsl1*, were also upregulated 5 and 7 days after Notch inhibition, and this upregulation was reduced in T3T4-treated mice (Figure S4N). Using transponders that we surgically implanted in BAT, we observed that aJ1/2 treatment increased BAT temperature in a manner that was largely rescued by T3T4 administration (Figure S1P). We next analyzed the levels of several metabolic substrates in serum and observed a reproducible decrease in glucose, triglycerides and NEFAs (Figures S4Q and S4R), consistent with increased BAT activity (Maliszewska and Kretowski, 2021). Treatment with exogenous T3T4 hormones rescued the levels of triglycerides and NEFAs but not those of other metabolites that were decreased, such as glucose or cholesterol, suggesting that Notch inhibition triggers other non-thyroid related metabolic defects (Figures S4Q and S4R). Finally, Oxymax indirect calorimetry showed that Jag blockade reduced energy expenditure, which is defined as the collective cost for maintaining homeostasis and physical activity, and this was restored after T3T4 administration (Figure 4I). Given that thermoregulation is one of several components of energy expenditure, thyroid-dependent decreases in energy expenditure are consistent with the thermoregulation defects observed upon Notch inhibition.

Altogether these data show that Notch inhibition, through decreases in T4 and T3 hormones, causes thermoregulation defects that mimic hypothyroidism in mice (Ueta et al., 2011; Warner et al., 2013; Weiner et al., 2016). We suggest that Notch inhibition increases heat dissipation that contributes to a decrease in body temperature that subsequently leads to compensatory BAT activation. Consistent with hypothyroidism as a root cause, exogenous administration of thyroid hormones rescues these thermoregulation defects.

### Genetic perturbation of Notch signaling induces hypothyroidism in mice and human

To directly test whether Notch signaling in thyrocytes contributes to the phenotypes described above, we used tamoxifen induction of thyroglobulin (Tg)-CreER^T2^ (Undeutsch et al., 2014) in Notch2^fx/fx^ mice to inducibly and specifically delete the Notch2 gene in thyrocytes (McCright et al., 2006). We focused on Notch2 because it is the dominant active receptor in thyrocytes (Figures 2A and S2A). Tamoxifen administration induced expression of Cre and efficient recombination at the Notch2 locus, which led to decreased Notch2 expression in thyrocytes (Figures S5A-C).

Following genetic deletion of N2 in thyrocytes and treatment with N1 blocking antibody, we observed reductions in Notch thyroid target gene expression, T4 serum levels, and body temperature, as well as increases in heat dissipation (Figures 5A-C and S5D-E). As observed following antibody-based Notch inhibition (Figure 4D and 4E), the decreased body temperature seen here likewise correlated with increased BAT activation, based on decreases in BAT weight, lipid droplets in BAT tissue, circulating triglyceride levels, and energy expenditure (Figures 5D-F and S5F). Importantly, these genetic data establish that thyrocyte N2 function is sufficient to explain the N2 contribution to thyroid dysfunction and the downstream thermoregulation defects.

**Figure 5.**
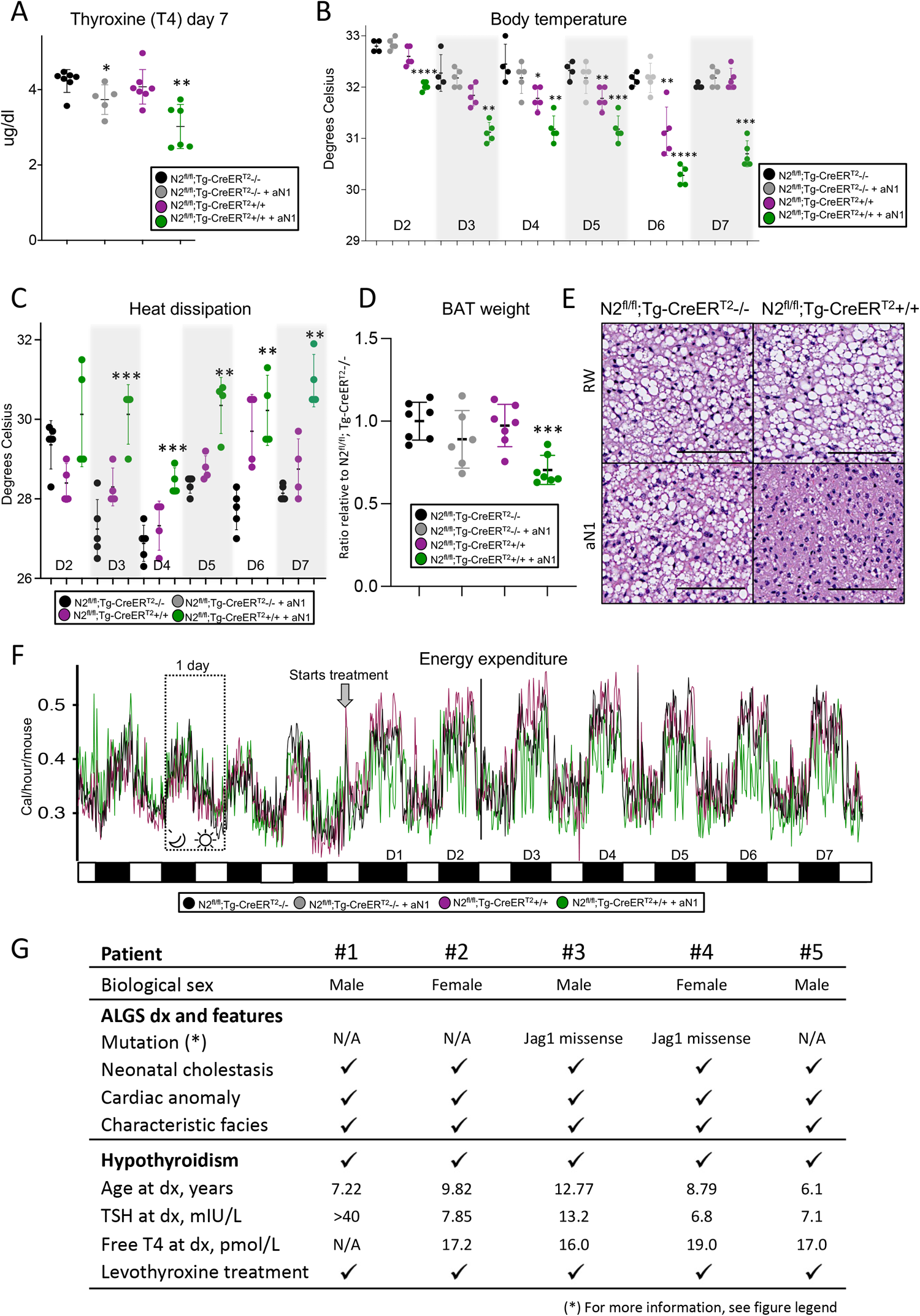
Genetic alterations in Notch induce hypothyroidism in mice and humans. **(A)** Serum levels of thyroxine (T4) in mice (n ≥ 5) of the indicated genetic strain and treatment, measured by ELISA at day 7. All mice were treated daily for 4 days with 200mg/kg of tamoxifen to genetically delete N2 in Tg-CreER^T2^+/+ mice. N2^flx/flx^;Tg-CreER^T2^ -/-treated with tamoxifen served as controls. At day 15, mice were injected with 5 mg/kg of either aRW (isotype control) or aN1. 2 independent experiments, Ave. ± s.d. **(B)** Body temperatures measured using a rectal probe at the same time on each of the indicated days in mice (n ≥ 6) housed at 22°C and treated as in (A). 2 independent experiments, Ave. ± s.d. **(C)** Infrared thermography measurements of heat dissipation from the tails of mice (n = 4) treated as in (A). Ave. ± s.d. **(D)** Wet weight of brown adipose tissue (BAT) in mice (n = 6) treated as in (A). 2 independent experiments, Ave. ± s.d. **(E)** Representative haematoxylin and eosin staining of brown adipose tissue (BAT) sections obtained after 7 days from mice (n = 5) treated as in (A). Scale bars = 100mm. **(F)** Energy expenditure in mice (n = 5) treated as in (A). Black and white bars represent daily night and day cycles, respectively. Oxygen and carbon dioxide levels were measured every 15 min for 3 days before and 7 days after treatment, as indicated. Ave. ± s.d. **(G)** Summary of data collected from Alagille syndrome patients diagnosed with hypothyroidism. Reference ranges: TSH 0.5-5.3 mIU/L; free T4 10-28 pmol/L. Levothyroxine is the manufactured form of T4, used as a standard of care to treat hypothyroidism. (*) Subject 3: c.752G>T missense mutation in exon 5 of Jag1, leading to a p.Cys251Phe protein change, variant classified as likely pathogenic. Subject 4: c.784T>C missense mutation in exon 6 of Jag1, leading to a p.Cys262Arg protein change, variant classified as pathogenic. Statistical significance was assessed using the unpaired two-tailed Student’s t-test with Welch’s correction: *p*<0.05, *; *p*<0.01, **; *p*<0.001, ***; *p*<0.0001, ****. See also Figure S5.

However, neither N2 genetic disruption nor N1 antibody blockade alone was sufficient to induce these phenotypes, consistent with our results that combined blockade of N1 and N2 is required to induce hypothyroidism (Figure S4B). We propose two models, not mutually exclusive, to explain this dual requirement for N1 and N2. First, N1 inhibition in a non-thyrocyte cell population may be required along with N2 inhibition in thyrocytes or, second, N1 activity in thyrocytes (typically below the level of our detection method, Figures S1H and S1I), may compensate for N2 loss. Our detection of N1 activity in a subset of thyrocytes in Notch2^fl/fl^; Tg-CreER^T2^ mice (Figure S5H) lends support to this second hypothesis.

While murine and human hypothyroidism both cause similar thermoregulation defects (Samuels et al., 2017), hypothyroidic mice lose weight whereas hypothyroidic humans gain weight (Kaspari et al., 2020). Such opposing phenotypes invite questions of whether murine thyroid biology accurately models such biology in humans and, specifically, whether our findings of Notch control of thyroid function are relevant in humans. We therefore analyzed data from a cohort of children with Alagille syndrome (ALGS), an autosomal dominant multisystemic disorder caused by loss-of-function mutations in JAG1 or NOTCH2. The ALGS phenotype is typically dominated by cholestatic liver disease with bile duct paucity, in association with an array of other clinical features that are variably expressed, including congenital cardiac defects, ocular and skeletal malformations, vascular and renal anomalies, and characteristic facial features (Kamath et al., 2018). Although isolated case reports take note of thyroid defects in ALGS (Filippis et al., 2016), thyroid biology in Alagille patients has not been carefully studied, and any possible clinical hypothyroidism seems to have been largely dismissed as an indirect and downstream consequence of the well-known liver problems. Among a cohort of 158 patients with genetically and/or clinically defined ALGS (age ranging from 0.4-22.82 years), we analyzed data from a subset of patients (n=72) in whom a thyroid history and T4 thyroid hormone serum levels had been assessed. We identified 5/72 (7%) patients with significantly lower T4 and increased TSH levels who were diagnosed with hypothyroidism, requiring thyroid hormone replacement therapy (Figure 5G), compared to the 0.1% estimated incidence of pediatric hypothyroidism overall (Hunter et al., 2000). Given that thyroid testing is not routinely performed in Alagille patients, this 7% versus 0.1% comparison likely underestimates the true enrichment of Alagille patients with thyroid defects. In any case, the detected enrichment provides a link between Notch signaling and thyroid function in humans and suggests that the biology we describe in our mouse models is at work in humans as well.

## DISCUSSION

Despite the essential role of the thyroid and the high prevalence of thyroid diseases, a fine-resolution view of the cellular composition of this crucial organ has been lacking. Although differences in thyrocyte morphology and growth were observed decades ago (Huber et al., 1990; Roger et al., 1992; Studer et al., 1989), their physiological significance has remained unknown and modern single-cell technologies had yet to be deployed to define cellular subpopulations and heterogeneity. Our single-cell and spatial transcriptomic analyses now address this gap in the murine thyroid for the first time and revise previous paradigms by revealing two major thyrocyte subpopulations instead of a single homogeneous population. Our two main types of murine thyrocytes, TFC1 and TFC3, differ in their metabolic activities, expression of classic thyrocyte lineage markers, and levels of Notch signaling. TFC1 cells not only appear highly metabolically active but express the full array of historically-defined thyrocyte markers. We suggest that TFC1 cells, therefore, best resemble classically defined thyrocytes. In contrast, TFC3 cells lack detectable expression of numerous thyrocyte markers but express progenitor markers and appear less active metabolically. We speculate that TFC3 cells may represent a relatively quiescent and less-differentiated/progenitor thyrocyte state, a compelling question to address in future research. Recent studies in zebrafish also identified two distinct thyrocyte subpopulations with similar expression and metabolic patterns as our TFC1 and TFC3, suggesting that these thyrocyte classifications may be conserved in vertebrates (Gillotay et al., 2020).

Notch has been classically described as an arbiter of cell fate during development, including in the thyroid, and dysregulation of these presumed developmental signals has been observed in thyroid cancer (Ferretti et al., 2008); however, our studies point to a novel role for Notch in the physiology of the normal adult thyroid. Our sequencing and pathway analyses led us to test the hypothesis that higher Notch signaling in TFC1 versus TFC3 cells might drive the differentiation-state differences that we uncovered in these thyrocyte subpopulations. Notch activity parallels expression of thyrocyte differentiation markers during thyroid development and cancer (Ferretti et al., 2008) and regulates Pax8 expression in other tissues (Jayasena et al., 2008; Kessler et al., 2015). We not only discovered similar parallels in TFC1 versus TFC3 cells but also showed that Notch inhibition decreased expression of the full thyrocyte lineage program in TFC1, including of Pax8. We thus conclude that Notch, directly or indirectly, controls such a lineage program in a subset of adult thyrocytes.

We also discovered that Notch controls the metabolic activity of adult thyrocyte mitochondria. A role for Notch in mitochondrial biology has been reported although not thoroughly characterized. For example, Notch has been described as: reprogramming mitochondrial activity in macrophages (Xu et al., 2015), supporting growth-promoting mitochondrial integrity in breast cancer cells (Landor et al., 2011), and regulating the mitochondrial proteome by increasing mitochondrial mass and membrane potential (Basak et al., 2014). We now establish, using a battery of analyses of the thyroid gland and a thyrocyte cell line, that Notch inhibition disrupts thyrocyte mitochondrial morphology and function, most notably decreasing the ROS production required for thyroid hormone synthesis. These mitochondria studies thus provide an explanation for the decreases in circulating levels of these hormones that we observe following Notch blockade in mice. Overall, we have revealed an unappreciated and essential role for Notch in thyroid hormone production and provided a root cellular mechanism. We speculate that the importance of such links between a fate-controlling pathway and mitochondria biology may carry broad implications beyond the thyroid; indeed, cellular and mitochondrial metabolism are emerging as potential drivers of cell fate changes (Döhla et al., 2022), raising the provocative possibility that Notch control of some fate decisions might be mechanistically linked to Notch regulation of mitochondrial activity.

Although some metabolic aspects of thyroid biology appear distinct between mice and humans, notably with hypothyroidism oppositely affecting body weight in each species, thyroid control of thermoregulation appears conserved (Iwen et al., 2018). We suggest that our discovery linking Notch signaling to thyroid hormones and thermoregulation impacts longstanding interpretations of drug responses and pathological mechanisms in patients. For example, pan-Notch inhibition in humans may induce many of the same thermoregulatory changes that we have observed in mice. Furthermore, our analysis of ALGS suggests that decreases in Notch signaling directly induce hypothyroidism in humans. ALGS is characterized by loss-of-function mutations in JAG1 and NOTCH2, a ligand-receptor pair that we found important for mouse thyroid function. Our mouse results suggest that the hypothyroidism we found enriched in Alagille patients reflects direct defects in Jag1-Notch2 signaling in the human thyroid and not, as appears to have been widely assumed, indirect consequences of Notch defects in the liver or other tissues. Our discovery thus carries important implications for clinical practice, suggesting that patients with ALGS are at higher risk for hypothyroidism and should be routinely screened for thyroid dysfunction as a primary symptom of decreased Notch signaling. In addition to ALGS, dysregulated Notch signaling has been linked to cancer, inflammation and metabolic disorders (Allen and Maillard, 2021; Aster et al., 2016; Xu and Wang, 2021), and thus researchers have developed numerous therapeutics to block Notch signaling, including pan-Notch antagonists such as gamma-secretase inhibitors (GSIs). Our discovery importantly predicts that hypothyroidism has likely been missed as an on-target toxicity in patients treated with such pan-Notch inhibitors. Indeed, a patient-reported side-effect of GSI treatment has been decreased energy (Krop et al., 2012), and we speculate that such reports may reflect blockade of Notch-controlled whole-body metabolism.

Finally, thyroid dysfunction significantly impacts human health, underlies the most prevalent endocrine disorder, and correlates with obesity, insulin resistance, hypertension, dyslipidemia and aging (Taylor et al., 2018). Clinical manifestations of hypothyroidism are highly variable, ranging from metabolic disorders, fatigue, cold intolerance, infertility and mental disorders and, when undiagnosed or untreated, can lead to severe complications such as heart failure, psychosis and coma (Roberts and Ladenson, 2004). Our discovery places thyrocyte Notch signaling at the apex of whole-body homeostasis, thermoregulation and metabolism, and thus has important implications for human health. Overall, our in-depth and cellularly detailed characterization of the thyroid generates novel insights into the normal function and pathological dysfunction of this key metabolic organ, with important implications for clinical practice, including in patients with ALGS.

## ACKNOWLEDGEMENTS

We are grateful to Kerstin Seidel, Charisa Cottonham and Patricia Himmels for experimental support and discussion, Mark Chen and Michael Stolzenberg for helping score brown adipose tissue activity, the Genentech Research Pathology and Next-Generation Sequencing labs and Brad Bolon and Dennis Wilson (GEMpath, Inc.) for their experimental contributions. We thank Shingo Kajimura, Antonio Zorzano and Navdeep Chandel for sharing methods and for expert feedback on metabolism and mitochondria results, and Ricardo Pujol and Daniel Alvarez for sharing their thyroid human data and feedback. We appreciate the insightful feedback and comments on the paper from Felipe de Sousa e Melo, Ciara Metcalfe, Weilan Ye and Heinrich Jasper.

## AUTHOR CONTRIBUTIONS

L.M designed and performed the experiments described below, analyzed and discussed data throughout the manuscript and co-wrote the manuscript. T.T.N and R.P analyzed the scRNA-seq and bulk RNAseq data, performed Topic analysis and discussed the data. S. H. provided experimental assistance with the genetic mouse model. M.R. performed the cryo-EM experiments and analysis. S.M.V and B.M.K collected the thyroid data from ALGS patients. Z.L, F.K.C and D.S performed the thyroid metabolomics experiment and analysis. A.S. provided histopathological analysis and discussion. C.W.S supervised the study, provided feedback on experimental design and data interpretation, and co-wrote the manuscript. All authors discussed the results and commented on the manuscript.

## MATERIALS AND METHODS

### Animal strains and treatments

Animal experimentation at Genentech was performed according to protocols approved by the Genentech Institutional Animal Care and Use Program (IACUC) committee. Male C57BL/6 mice were obtained from Jackson Laboratories (strain ID: 00664). Ob/Ob mice were obtained from Jackson (strain ID: 00632). N2fl/fl (Jackson #010525) were crossed with Tg-Cre-ERT2 mice (Jackson #030676) to generate N2fl/fl;Tg-Cre-ERT2 mice. All mice were housed under specific pathogen-free (SPF) conditions and only male 10-14 weeks of age were used. Mice were injected intraperitoneally with a single dose of blocking antibodies diluted in PBS at the following concentrations: anti-Jag1, 20 mg/kg; anti-Jag2, 20 mg/kg; anti-Notch1, 5 mg/kg, anti-Notch2, 10 mg/kg; anti-Notch3, 20 mg/kg (injected twice a week). Anti-Ragweed isotype control antibody was injected at concentrations to achieve equal dose of the combination of antibodies in each study. Exogenous administration of thyroxine (T4) and 3-Iodotrionine (T3) was performed by intraperitoneal injection of the hormones diluted in PBS from a stock solution prepared fresh. Stock solution of T4 was prepared at 0.2μg/ml in 0.01M NaOH, 0.1% BSA in PBS and stock solution of T3 was prepared at 1ug/ml in 4mM NaOH in PBS. The administration of the combination of T3+T4 was performed daily starting at day 2 of the antibody treatment with a dose 1ug/g BW of T4 and 0.05μg/g BW T3. Tamoxifen was diluted in corn oil at 40mg/ml and administered to the mice by oral gavage at 5ml/g BW to achieve a 200 mg/kg. Tamoxifen administration was performed daily during 4 days, followed by 3 days resting period.

### Genotyping and recombination PCR

DNA from N2fl/fl;Tg-Cre-ERT2 mice was amplified using the following primers 5’-CAA CCC CAG ATA GGA AGC AG-3’ and 5’-GAG CCT TTT CCC CAT ATT CC-3’ and PCR program 94°C/4 min, (94°C /1 min, 60°C /30 sec, 72°C /1 min) 30 cycles, 72°C /10 min. PCR products sizes were 202bp for the WT allele and 285bp for the transgenic allele. DNA from N2fl/fl;Tg-Cre-ERT2 thyroids was extracted using All prep DNA/RNA extraction kit (Qiagen, 80284) and DNA was amplified by PCR using primers 5’-GAG CCT TTT CCC CAT ATT CC-3’ and 5’-GCTCAGCTAGAGTGTTGTTCTTG-3’ and PCR program 94°C/4 min, (94°C /1 min, 60°C /30 sec, 72°C /2.5 min) 30 cycles, 72°C /10 min. PCR products sizes were 1485bp for the WT allele and 450bp for the recombined allele.

### Thermoregulation studies

Body temperature was measured using a rectal fiber optic gauge (Fiso, FTI-10). BAT temperature was measured by IPT300 (Implantable Programable Transponders), implanted with a 15G trocar in the interscapular area. Telemetry studies were performed using DST nanoRF-T transponders (Star Oddi), implanted in the abdominal cavity of mice. For that, mice were anesthetized with 4% isoflurane, and the previously sterilized implants were introduced by a small incision in the abdominal cavity. Slow released anesthetic (Buprenorfine) was administered at the time of the surgery and the antibody treatments were performed 30 days after the surgery. The temperature data was collected using Mercury and Gná software (Star Oddi). For infrared thermography, pictures of mice were taken using the FLIR T4xx series infrared camera. For cold challenge, we used the Columbus cabinets (IT79SD) and mice were acclimatized to 4°C or 30°C for a week prior to the antibody treatments. Measurements of oxygen consumption and carbon dioxide production were taken in Oxymax indirect calorimeter chambers (Columbus Instruments, 120455 and 110113). Measurements were collected using Clams Comprehensive Lab Animal monitoring system and analyzed using CLAMS data eXamination (CLAX) software.

### Serum analysis

Blood was collected at the time of necropsy in EDTA-treated tubes and centrifuged for 10min at 10g to isolate the serum. Serum biochemistry was analyzed by Beckman AU480 using Beckman Coulter kits: Gluc (OSR6221); Chol (OSR6216), Trig (OSR61118) and NEFAs (Randox, FA115). The analysis of hormone levels in serums was performed with ELISA kits following the manufacturer protocol: T4 (IBL America, IB19133), T3 (MSB727732), TSH (GBiosciences, IT6045), TRH (TSZ, M7491) and Calcitonin (LSBio, LS-F23047-1).

### Histopathological analysis and stainings

Tissue samples were collected at the time of necropsy and fixed in in 10% neutral buffered formalin (4% formaldehyde in solution), paraffin-embedded, and cut in 4 μm sections, which were mounted in superfrost®plus slides and dried. Serial sections were stained with hematoxylin and eosin (HE) or Alcian Blue (Millipore, TMS010C) followed by nuclear fast red counterstaining (Vector, H3403). Thyroid morphology (assessed by 3 independent pathologists: Alexis Scherl, Brad Bolon and Dennis Wilson) was evaluated for peripheral follicle size, follicular epithelial cell height, and relative density of colloid resorption vacuoles. Follicles with attenuated epithelium and less than 4 colloid resorption vacuoles were considered inactive. Thyroid sections estimated to have more than 40% of peripheral follicles inactive were interpreted to have reduced activity. Scoring based on epithelial features resulted in flattening epithelium in 10-40% of aJ12 and >80% in aN123 (and aN12). The scoring criteria used for the thyroid gland were adapted from published descriptions of function-dependent changes in mice of various ages (Lee et al., 2016).

Immunohistochemistry against Nkx2-1 (ab76013) and cleaved (active) NICD1* (Cell Signaling, 4147) was performed using an automated immunostaining platform (Ventana discovery XT, Roche). Antigen retrieval was first performed with high pH buffer (CC1, Roche), endogenous peroxidase was blocked and slides were then incubated with the appropriate primary antibodies as detailed: Nkx2-1 (1:100) and NICD1* (1:100). After the primary antibody, slides were incubated with the corresponding secondary antibodies and visualization systems (OmniMap Rabbit HRP, Ventana, Roche) and counterstained with hematoxylin (Ventana, Roche). Finally, the slides were dehydrated, cleared and mounted with a permanent mounting medium (Sakura, 6419) for microscopic evaluation. Immunohistochemistry against Pax8, Calcitonin, Calcitonin gene-related protein, N2, UCP1 and Cre was performed manually using standard techniques. In brief, sections were deparaffinized in xylene, rehydrated in ethanol and antigen retrieval was performed in DAKO target retrieval solution (#2369) for 15min. Endogenous peroxidase was blocked with 3% H2O2 for 20min and slides were blocked in 5%BSA in PBS 0.1% Triton for 1 hour. Overnight incubation at room temperature (RT) was performed with the appropriate primary antibodies diluted in blocking buffer as follows: Pax8 (1:100, CS9857S), Calcitonin (1:100, PP072AA), Calcitonin gene-related peptide (1:100, ab47027), N2 (1:500, CS5732S), UCP1 (1:100, ab23841) and Cre (1:100, CS15036S). Slides were incubated with Secondary antibody powervision poly-HRP anti-rabbit IgG (Leica PV6119) for 1 hour, and developed with 30-diaminobenzidine tetrahydrochloride (DAB). Counterstaining with hematoxylin (1:10) was performed and slides were dehydrated with gradual ethanol and xylenes. Finally, slides were mounted with Permount for microscopic evaluation. RNAscope in situ hybridization was performed manually using RNASscope chromogenic brown assay (ACD, 322310) with the following probes: Jag1, 412831; Jag2, 417511; Notch1, 404641; Notch2, 425161; Notch3, 425171 and Nrarp, 411771 and following the manufacturer recommendations. RNAscope for HeyL, Hey1 and Nrarp was performed and quantified by ACD. Cox4 staining (VitroView™, VB-3022) was performed in thyroid cryosections following manufacturer’s recommendations.

Double immunofluorescence against NICD1* and Pax8, NICD1* and calcitonin and NICD1* and endomucin was performed similarly to manual immunohistochemistry until the overnight incubation with NICD1* (CS4147S), that was performed at room temperature (RT) and 1:500. Slides were incubated with Secondary antibody powervision poly-HRP anti-rabbit IgG (Leica PV6119) for 1 hour. TSA amplification (Perkin-Elmer) was performed with Cy3 (1:100) for 30min. Slides were then blocked again in 5% BSA in PBS-Tween PBS 0.1% Triton for 1 hour. Overnight incubation at room temperature (RT) was performed with the appropriate primary antibodies diluted in blocking buffer as following: Pax8 (ab53490, 1:100), Calcitonin (NBP1-46397, 1:100) and endomucin (ab106100, 1:100). Slides were incubated with the corresponding secondary fluorescent antibody (1:250) for 1h followed by 15min DAPI staining. Finally, slides were mounted with Prolong Gold antifade reagent (P36934) for microscopic evaluation. Immunofluorescence against cleaved (active) NICD2* (40-2-7, PUR81369) was performed similarly to the double immunofluorescence (NICD2* incubated overnight at 1:500, followed by TSA amplification) until after TSA amplification. Slides were then incubated with the corresponding secondary fluorescent antibody (1:250) for 1h followed by 15min DAPI staining. Finally, slides were mounted with Prolong Gold antifade reagent (P36934) for microscopic evaluation. Immunofluorescence against Pax8 (ab53490, 1:100), Tpo (ab133322) and Slc5a5 (NBP2-33547) was performed similarly, without TSA amplification, and incubated with the corresponding secondary antibody (1:250) for 1h followed by 15 min DAPI staining. Immunofluorescent images were taken using Leica DMi8 Thunder microscope and quantifications were done using QuPath 0.2.3 software.

### Thyroid tissue processing and imaging with BSE-SEM

Thyroid tissue was fixed by immersion fixation in modified Karnovsky’s fixative (2.5% paraformaldehyde and 2% glutaraldehyde in 0.1 M cacodylate buffer, pH 7.2) for at least 24 hours at 4°C (Karnovsky, 1965). Fixed tissue was washed with ultrapure water and post-fixed with 2% reduced osmium tetroxide for 2 hours. The samples were then washed again in ultrapure water and stained “en block” with 0.5% (w/v) uranyl acetate at 4°C overnight. Following staining, samples were dehydrated in a series of ascending acetone concentrations, rinsed twice with finally embedded in epoxy resin Eponate-12 (Ted Pella, Redding, CA, USA). Semithin sections (500 nm thickness) were cut with the UMC ultramicrotome (Leica Biosystems, Buffalo Grove, IL, USA) using a DIATOME diamond knife for histology (Electron Microscopy Sciences, Hatfield, PA, USA). Sections were transferred to carbon-coated histology glass slides and dried. Finally, sections were stained with 4% aqueous uranyl acetate for 15 min and 0.1% Reynold’s lead citrate for 1 min to enhance contrast (Reynolds, 1963). Sections were thoroughly rinsed with ultrapure water and dried on a heat plate before being transferred to the SEM. Scanning electron microscopy (SEM) was performed using a GeminiSEM 300 equipped with a field emission gun (Carl Zeiss AG, Oberkochen, Germany). For operation of the GeminiSEM 300 microscope the application software SmartSEM (version 6.01) was used (Carl Zeiss AG, Oberkochen, Germany). Imaging was with the backscatter electron detector (BSD1) at 8.5 mm working distance, 60 μm aperture, 5keV acceleration voltage and with operation of the field emission gun in “high current” mode. Image size was at least 4096 × 3072 (4k x 3k) pixels. For imaging of ultrastructural detail pixel sizes between 1-3 nm, and magnifications from 10.000x to 25.000x were used. The greyscale of the images was inverted to achieve TEM-like representations.

### Thyroid single cell isolation

Thyroid tissue was extracted from mice at the time of necropsy. To generate single cells, thyroids were cut with a razor blade and digested for 30min at 42°C. The digestion was done with 0.05mg/ml Liberase TM (Sigma, 5401127001) diluted in Advanced DMEM/F12, 10mM HEPES, 1% Glutamax, 1% P/S and 1:1000 Rock inhibitor (Y27632, StemCell 72304). After digestion, centrifuge the cells at 1200rpm/5min, wash them with Advanced DMEM/F12, 10mM HEPES, 1% Glutamax, 1% P/S and filter them through a 70um strainer. This single cell isolation will be used for organoids formation, single cell RNAseq and mitochondrial stainings for FACs (TMRM, mitosox, mitotracker and DCFD).

### Thyroid organoids isolation, culture and characterization

Single cells were isolated from the thyroid (see *Thyroid single cell isolation*). A pool of 7 thyroids was digested into single cells in one experiment (45ml of digestion) while pools of 2 thyroids (12ml digestion) were digested in the other experiment. Count cells and plate them into 50μl Matrigel (Corning, 356255) plugs (3 plugs in the first experiment and 1 plug per digestion in the second) and culture with thyroid media. Thyroid culture media is Advanced DMEM/F12, 10mM HEPES, 1% Glutamax, 1% P/S, 1:50 B27, 2.5mM (ThermoFisher, 17504044), 2.5mM N-Acetylcisteine (Sigma A7250), 50ng/ml mr-EGF (peprotech 315-09), 10μg/ml hrFGF10 (Peprotech, 100-26), 16mIU/ml TSH (Lee Biosolutions, 996-51), 50ng/ml R-Spondin (Peprotech 120-38), 25ng/ml hrNoggin (Peprotech 250-38), and 100ng/ml Wnt3a (Peprotech 315-20).

For immunofluorescence staining, plate thyroid organoids in 8-well chamberslides (plugs of 30μl Matrigel). After 3-4 days of plating, fix the organoids with 4% para-formaldehide (PFA) for 20 min. Wash with PBS and permeabilize with 0.5% Triton x100 in PBS for 30 min. Block for 1 h in PBS+ 0.5% Triton x100 +1% BSA +10% FBS +0.4% Tween 20 and incubate overnight with primary antibodies at 4°C. The primary antibodies and dilution used were Pax8 (ab53490, 1:100), Calcitonin (NBP1-46397, 1:100) and Nkx2-1 (ab76013, 1:100). Wash and stain with DAPI for 30 min. Take slides out of the chambers and mount them with Prolong Gold antifade reagent (P36934) for microscopic evaluation. Imaging was done with the SP8 invested confocal microscope.

### FRTL5 culture, staining and Seahorse assay

FRTL5 was purchased from Cell Line Services (CLS) and cultured FRTL5 media (CLS): Coon modified Ham’s F12 medium supplemented with L-glutamine, 10μg/ml insulin, 10nM hydrocortisone, 5μg/ml transferrin, 10ng/ml somatostatin, 10ng/ml glycyl-L-histidyl-L-lysine acetate, 20ng/ml TSH and 5% bovine calf serum. To expand the cells, use accutase (Stem Cell, 07920) for 10 min at RT and centrifuge 5 min at 300g. For immunofluorescence, plate 50K cells in chamberslides (LabTek, 154534) and fix with 4% PFA for 20 min. Images were taken using Leica DMi8 Thunder microscope. For Seahorse assays, plate 15K in Seahorse microplates (Agilent, 101084-004) with FRTL5 media, add antibodies (25μg/ml) or the GSI Compound E (Cmp. E) (1μM) the following day. Perform the Seahorse assays 2 days after the antibodies treatment in Seahorse xFe96 Analyzer reader using the Seahorse kits (Cell Mito Stress: 103015-100 or Real time ATP: 103592-100) following manufacturer’s recommendations. Results were normalized to cell counts using BioTek Cytation 1 image reader and Hoeschst 33342 staining (2μM/well).

### TMRM, mitosox, mitotracker and DCFDA stainings and analysis

Thyrocyte were isolated from thyroids of mice that had been treated *in vivo* with the antibodies for 7 days (see *Thyroid single cell isolation*). Cell were incubated at 37°C for 1h in thyroid organoid media with mitotracker green 1:1000 (Invitrogen, M7514) and mitosox red 1:500 (Invitrogen, M36008) or in PBS with TMRM 1:1000 (ImageIT, I34361), followed by 1:1000 DCFDA/H2DCFDA (ab113851). For controls THBP 1:250 (ab8206006) or 10mM N-acetylcisteine (NAC, A7250) were used. Fluorescence intensity was evaluated by FACSymphony A3 Dual and analyzed by FlowJo.

### Cox4, GSH and catalase enzymatic assays

Thyrocyte were isolated from thyroids of mice that had been treated *in vivo* with the antibodies for 7 days. Thyroid tissues were manually disrupted using a Dounce homogenizer. Enzymatic activities were determined using the following kits and the recommended protocols: Complex IV (ab109911), catalase (ab83464), GSH/GSSG (ab205811). For GSH/GSSG deproteinization was performed using a deproteinization kit (ab204708)

### RNA isolation, qRT-PCR and RNAseq

Total RNA was extracted from BAT and thyroid tissues with Trizol (Invitrogen) following the provider’s recommendations. After adding isopropanol, the samples were transferred into RNAeasy extraction column (Qiagen), washed and eluded following the manufacturer protocol. For Taqman, RNA was retrotranscribed and amplified in an QuantStudio 7 Flex thermocycler (Applied Biosystem) using Taqman RNA-to-Ct 1-Step Kit (Thermofisher, 4392653). The Taqman probes used were the following Jag1 (Mm00496902_m1), Notch1 (Mm00435249_m1), Hes1 (Mm01342805_m1), Nrarp (Mm00482529_s1), Hey1 (Mm00468865_m1), Heyl (Mm00516555_m1), Myc (Mm00487804_m1), Nkx2-1 (Mm00447558_m1), Foxe1 (Mm00845374_s1), Pax8 (Mm00440623_m1), Slc5a5 (Mm01351811_m1), Tshr (Mm00442027_m1), Tpo (Mm00456355_m1), Trp63 (Mm00495793_m1), Calca (Mm00801463_g1), Ascl1 (Mm03058063_m1), Bax (Mm00432051_m1), Atg3 (Mm00471287_m1), Il6 (Mm00446190_m1), actb (Mm02619580_g1).

The libraries were sequenced on Illumina HiSeq4000 (Illumina). We obtained on average 40 million single-end RNA-seq reads (50bp) per sample. Sequencing reads were aligned to the mouse genome (NCBI Build 38) using GSNAP (Wu and Nacu, 2010) version ‘2013-10-10’, allowing a maximum of two mismatches per 50 base pair sequence (parameters: ‘-M 2 -n 10 -B 2 -i 1 -N 1 - w 200000 -E 1 --pairmax-rna=200000 --clip-overlap’). Lastly, gene expression levels were quantified by calculating the number of reads mapped to the exons of each RefSeq gene using the HTSeqGenie R package. Transcript annotation was based on the RefSeq database NCBI Annotation Releases 104 for mouse. Read counts were scaled by library size, quantile normalized and precision weights calculated using the “voom” R package (Law et al., 2014). Subsequently, differential expression analysis on the normalized count data was performed using the “limma” R package (Ritchie et al., 2015) by contrasting treated samples with control samples, respectively. Gene expression was considered significantly different across groups if we observed an |log2-fold change| ≥ 1 (estimated from the model coefficients) associated with an FDR adjusted P-value ≤ 0.05. In addition, gene expression was obtained in the form of normalized Reads Per Kilobase gene model per Million total reads (nRPKM) as described previously (Srinivasan et al., 2016).

### Single cell RNAseq and analysis

Single cells were isolated from the thyroid (*see Thyroid single cell isolation*). Given the low number of cells obtained from a single mouse thyroid, the thyroids from six mice were pooled into one sample. Three samples from each treatment (aRW, aJ12, aN12) were sequenced. After digestion and wash, cells were centrifuged at 1200rpm/5min and resuspended in PBS 3% FBS. Samples were processed for single-cell RNA-seq (scRNAseq) as described previously (Long et al., 2019) using the Chromium Single Cell 3’ Library and Gel bead kit v2, following the manufacturer’s manual (CG00052 Chromium Single Cell 3 Reagent Kits v2 User Guide RevA; 10x Genomics). Cell density and viability of the single-cell suspensions were determined by Vi-CELL XR cell counter (Beckman Coulter). All of the processed samples had more than 85% viable cells. Cell density was used to impute the volume of single-cell suspension needed in the RT master mix, aiming to achieve ∼8000 cells per sample. cDNAs and libraries were prepared following the manufacturer’s manual (10X Genomics). Libraries were profiled by Bioanalyzer High Sensitivity DNA kit (Agilent Technologies) and quantified using Kapa Library Quantification Kit (Kapa Biosystems, Wilmington, MA). Each library was sequenced in one lane of HiSeq4000 (Illumina) following the manufacturer’s sequencing specification (10X Genomics).

Sequencing data were processed as follows: Sample demultiplexing, barcode processing and single cell 3’ gene counting were performed using 10X Cell Ranger. Only cells with a mitochondrial UMI fraction <0.25 were considered for analysis. Per-gene UMI counts were normalized by the total number of transcripts per cell, scaled by the median number of transcripts across all cells and log-transformed for downstream analysis. Dataset alignment (between 1st and 2nd processing dates), cell clustering, visualization and differential expression analysis were performed as prescribed in the OSCA book (https://osca.bioconductor.org) (Amezquita et al., 2019). The pipeline merged together single-cell data generated from multiple batches. For each batch, it ran the quality control, normalization and variance modelling steps, and then applied MNN correction to merge all batches into a single coordinate space. Dimensionality reduction and clustering on the merged data were performed to obtain a common grouping for all cells. The analysis used the top 20 principal components for clustering and visualization, the 50 nearest neighbors (k=50) for graph-based clustering and performed marker detection using the AUC with a log-fold change threshold of 1. We tested for the enrichment of particular gene categories to identify relevant biological processes associated with (1) differential expression or (2) transcriptional programs (topics), by using functions from the “clusterProfiler” R package (Wu et al., 2021) and the MSigDB gene set collection (Liberzon et al., 2011). In the case of differential expression, treated samples were contrasted with control samples, genes were ranked based on the π statistic (π=logFC * −log10(p-val)) (Xiao et al., 2014) and subsequently gene set enrichment was performed using the clusterProfiler::GSEA function. In the case of transcriptional programs, we extracted the top 50 genes associated with specific topics and performed a hypergeometric test using the clusterProfiler::enricher function. To characterize cells based on the activity of transcriptional programs, we performed topic modeling using functions from the CountClust R package (Dey et al., 2017). We evaluated a range between 2 to 15 topics for their fit using the FitGoM function. A total of 7 topics were selected to represent the data after inspecting the AIC and BIC statistics. Top genes associated with each topic were identified by applying the ExtractTopFeatures function, which uses the relative gene expression profile of the GoM clusters and applies a KL-divergence based method to obtain a list of top features that drive each of the clusters.

### Spatial transcriptomics (Visium) and analysis

Thyroid tissue was embedded in OCT, flash frozen, cryosectioned at 10μm and stained with HE for tissue visualization. 5-6 cryosections were also collected for RNA extraction using RNAeasy Micro kit (Qiagen, 74004). Quality of the RNAs was determined by Tapestation (Agilent, 5067-5576 and 5067-5576) and samples with RIN>7 were selected. Tissue optimization (10X Genomics, 1000192) was performed following manufacturer’s protocol and determined that proper permeabilization of the thyroid was achieved within 3 min. The spatial transcriptomics assay was performed using Gene expression (10X Genomics, 1000186), Library construction (10X Genomics, 1000190) and dual index (10X Genomics, 1000215) kits and following manufacturer’s recommendations. Visium Spatial gene expression data were processed using 10X Space Ranger. For Visium data, dataset alignment (between multiple treatment conditions), cell clustering, visualization, calculation of gene set scores and differential expression analysis were performed according to best practices using Seurat (v3.0.0.9000) (Stuart et al., 2018).

### Thyroid metabolomics

All metabolite extraction procedures were kept on ice. Briefly, 350μL of cold methanol containing in-house metabolomics recovery stable isotope labeled internal standard and 200μL of cold chloroform were added to each thyroid tissue sample (17 - 26 mg). Samples were vortexed for 10 seconds, homogenized with beads for 2 minutes, and centrifuged at 4000 RPM for 5 minutes. 200μL of cold H_2_O were added to supernatants to perform liquid-liquid phase separation. 120μL of the aqueous layer were used for global metabolic analysis and 40μL were derivatized as for targeted analysis. Metabolomics data were acquired using two complementary LC-MS analytical assays. For global metabolomics, ACQUITY UPLC BEH Amide Column (2.1mm × 150mm × 1.7μm, 130Å, Waters Corporation, Milford, MA) was used to separate metabolites with mobile phase A of 100% H_2_O containing 10mM ammonium formate and 0.125% formic acid, and mobile phase B of 95% acetonitrile in H_2_O containing 10mM ammonium formate and 0.125% formic acid. For targeted metabolomics, a derivatization assay targeting TCA metabolites and other small organic acids including short chain fatty acids (SCFAs) was performed with method details previously reported (Jaochico et al., 2019; Tan et al., 2014). Data acquisition was achieved on a Shimadzu Nexera HPLC series system (Shimadzu, Kyoto, Japan) couple with a Thermo Q Exactive Plus Hybrid Quadrupole-Orbitrap Mass Spectrometer (Thermo Fisher Scientific, CA, USA) for global metabolomics analysis and Shimadzu Nexera HPLC series system connected to triple quadrupole mass spectrometer with linear ion trap (QTRAP 6500®, AB Sciex Instruments, CA, USA) for targeted metabolomics analysis. Injection volume of 3μL was used for sample analysis under Heated Electrospray Ionization (HESI) condition. Samples were run for both positive and negative modes for BEH Amide assay, and positive only mode for derivatization assay. The Q Exactive Plus Mass Spectrometer was operated with the following parameters: Sheath gas flow rate, 50 units; Aux gas flow rate, 13 units; Aux gas temperature, 425 °C; Capillary temperature, 263 °C; Spray voltage, 3500V for pos and −2500V for neg; Scan mode, Full MS scan with data-dependent MSMS acquisition. In Full MS scan, scan range is 60-900 m/z; resolution is 70,000; AGC target, 1×e^6; Maximum IT, 200 ms. In ddMS2 scan, top 5 ions are selected with isolation window of 1.5 m/z; resolution is 17,500; AGC target, 5×e^4; Maximum IT, 20 ms. Data processing software Chromeleon 7.3 (Thermo Fisher Scientific, CA, USA), and Polly platform (Elucidata Corporation, Cambridge, MA, USA) were used for metabolite identification, peak picking / peak integration, and statistical analysis, respectively. Metabolites were identified at Level 1 confidence (Dunn et al., 2013) by matching at least two independent orthogonal experimental data (accurate mass, isotopic ratio, retention time, and MSMS fragmentation pattern) against in-house compound retention time (RT) library. Global metabolomics batch trend analysis of stable isotope labeled internal standards and matrix Pool QC (quality control) samples were examined for %relative standard deviation (%RSD less than 12%) to validate system suitability and data robustness. For global metabolomics, relative quantification was obtained through either MS peak area of analyte or MS peak area ratio of analyte / internal standard. For targeted metabolomics absolute quantification (concentration) was obtained using MultiQuant software (AB Sciex Instruments, CA, USA) using surrogate matrix calibration curves of every analyte in the targeted panel (TCA cycle metabolites & small carboxylic acid metabolites) and the data was reported as ng/g of tissue.

### ALGS patients’ data collection

Children with a clinically and/or genetically confirmed ALGS diagnosis seen at The Hospital for Sick Children, Toronto, Ontario and The Children’s Hospital of Philadelphia, Philadelphia, Pennsylvania, between Jan-1980 and Aug-2021 were retrospectively reviewed for a co-diagnosis of hypothyroidism. A diagnosis of ALGS was made in accordance with standard clinical criteria, and clinical data were collected through electronic medical records. The following data points were collected: biological sex, affected ALGS gene (JAG1/NOTCH2) and associated variant details (exon/intron, DNA variant, protein, protein change), ACMG/AMP variant classification and predicted Coding effect, ALGS clinical features (liver, cardiac, facies, ocular, skeletal, renal, and vascular involvement), age at hypothyroidism diagnosis, thyroid function test results (thyroid stimulating hormone (TSH) and free thyroxine (free T4) and current treatment regimen. Variants were classified according to the American College of Medical Genetics and Genomics (ACMG) guidelines(Richards 2015). The study was approved by the institutional review boards (IRB) at each center, and a waiver of consent was granted by the respective IRBs.

### Data availability

The primary bulk, single cell and Spatial sequencing data have been deposited in the Gene Expression Omnibus repository (GSE200268).

