## Supplemental Information and Figures for "Discovery of thyrocyte heterogeneity reveals an essential role of Notch signaling in thyroid function and mammalian homeostasis"

**Figure S1. The thyroid is composed of a heterogeneous population of thyrocyte populations with distinct Notch activity profiles**

(A) Heatmap of cell types. Annotations were generated using the "AUCell" R package, which scores the enrichment of the curated gene set for a particular cell type among the expressed genes in each cell. A cell then has enrichment scores for all the cell types and will be assigned to the cell type that has the highest score. This heatmap annotates the number of cells assigned to a particular cell type (x-axis) per cluster (y-axis), revealing the cluster identity. n = 3 samples.
(B) t-distributed stochastic neighbor embedding (tSNE) plot for the TFC1, TFC2 and TFC thyrocyte clusters showing in blue the cells expressing the indicated markers, n = 3 samples.
(C) Visium spatial transcriptomic analysis showing the enrichment in the expression of the indicated markers in thyroid sections of control mice, n =2 thyroids. Heatmap legends indicate the topic score per well. Uniform Manifold Approximation and Projection (UMAP) of the spatial transcriptomic data showing 6 clusters. UMAP showing in blue the barcoded spots expressing the indicated markers. Heatmap legends represent normalized counts.
(D) Quantification of Pax8+ follicles in the two lobes of the thyroid, detected as in panel (E), suggesting possible asymmetric distribution of positivity. Follicles with at least 4 Pax8+ cells were counted as positive. Ave. ± s.d.; n $\geq$ 4 mice.
(E) Representative immuno-fluorescent and -histochemical staining images of Pax8 and Nkx2-1 in thyroid sections, revealing heterogenous levels of protein expression. 40X objective; scale bars = 20mm; n = 5 mice.
(F) Heatmap of the cluster-specific markers in TFC1, TFC2 and TFC3. n = 3 samples.
(G) tSNE plots for the TFC1-3 clusters showing that expression of the indicated markers is enriched in TFC1. n =3 samples.
(H) tSNE plots for the TFC1-3 clusters showing that expression of the indicated Notch pathway genes is enriched in TFC1. n = 3 samples.
(I) Expression of Jag1, Jag2, Notch1 and Notch2 mRNAs detected by RNAscope (amplified *in situ* hybridization) in control thyroid sections, n = 3 mice. Scale bars = 100mm.
(J) Representative double immunofluorescent detection in thyroid sections of gamma-secretase-cleaved (active) NICD1* plus Pax8 or NICD1* plus calcitonin. n = 5 mice. The lack of NICD1* co-staining with either marker indicates that, at least under these conditions, Notch1 activity is not derived from a thyrocyte or parafollicular cell. 40X objective; n = 5 mice.
(K) Representative images of immuno-histochemical or -fluorescent detection of NICD1*, NICD1* plus the endothelial cell marker endomucin, NICD2, and cleaved (active) NICD2*. 40X objective; scale bars = 100mm; n = 5 mice.

See also Figure 1.

Figure S1

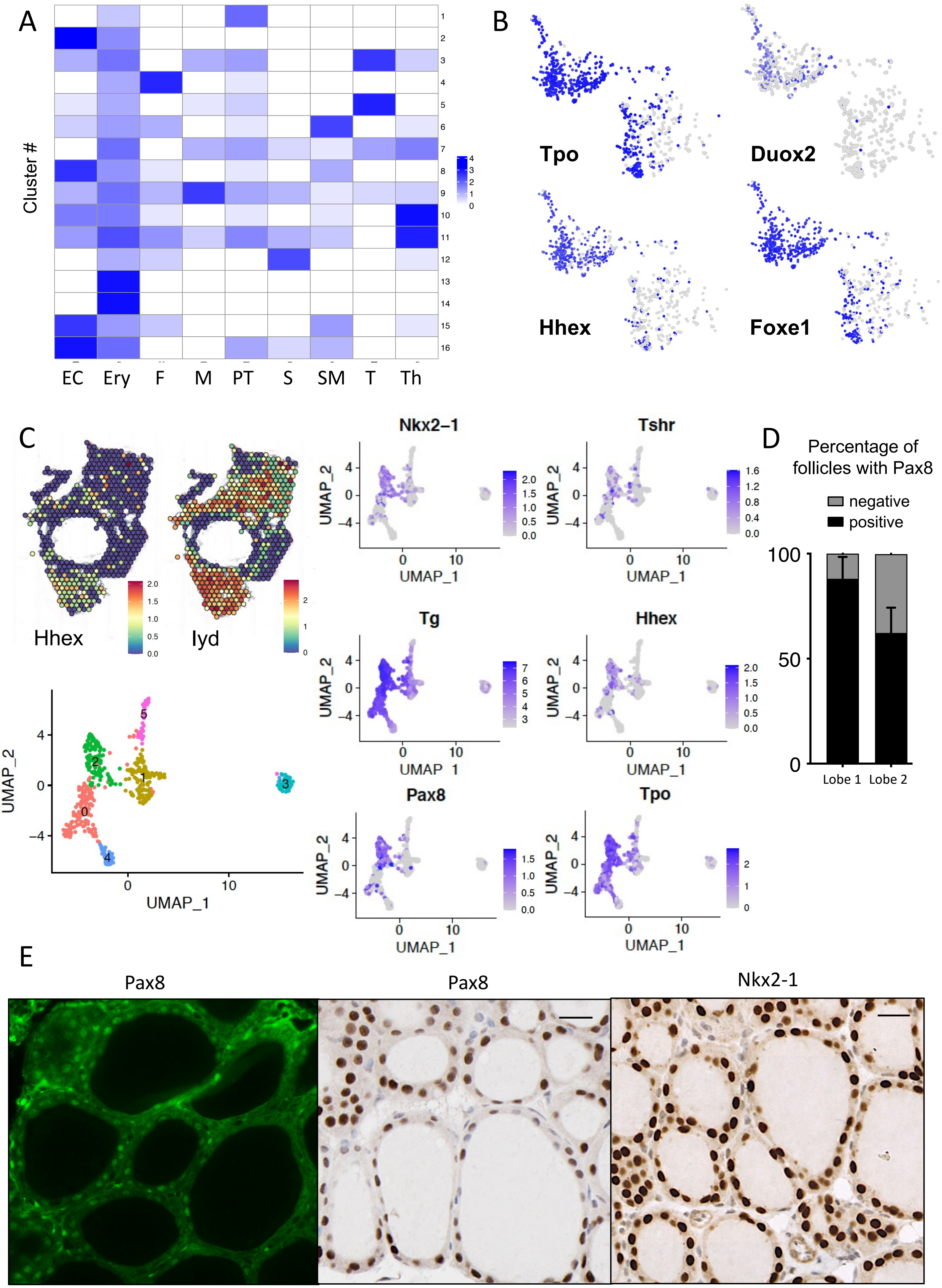
A
Cluster #
EC
Ery
F
M
PT
S
SM
T
Th
B
Tpo
Duox2
Hhex
Foxe1
C
Hhex
Iyd
Nkx2–1
Tshr
Tg
Hhex
Pax8
Tpo
UMAP_1
UMAP_2
D
Percentage of follicles with Pax8
negative
positive
Lobe 1
Lobe 2
E
Pax8
Pax8
Nkx2-1

Figure S1

F

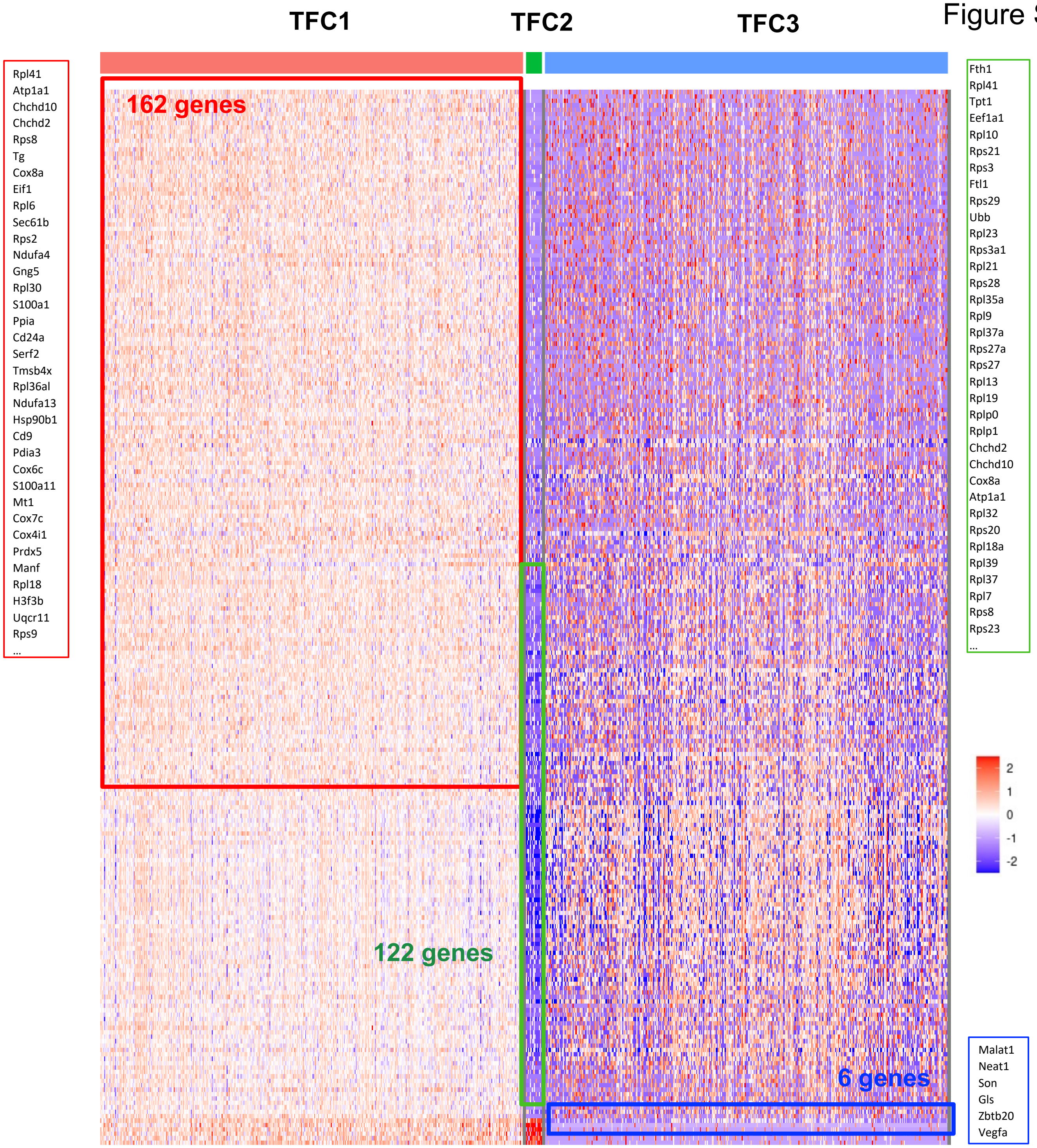
TFC1
TFC2
TFC3
162 genes
122 genes
6 genes
Rpl41
Atp1a1
Chchd10
Chchd2
Rps8
Tg
Cox8a
Eif1
Rpl6
Sec61b
Rps2
Ndufa4
Gng5
Rpl30
S100a1
Ppia
Cd24a
Serf2
Tmsb4x
Rpl36al
Ndufa13
Hsp90b1
Cd9
Pdia3
Cox6c
S100a11
Mt1
Cox7c
Cox4i1
Prdx5
Manf
Rpl18
H3f3b
Uqcr11
Rps9
...
Fth1
Rpl41
Tpt1
Eef1a1
Rpl10
Rps21
Rps3
Ftl1
Rps29
Ubb
Rpl23
Rps3a1
Rpl21
Rps28
Rpl35a
Rpl9
Rpl37a
Rps27a
Rps27
Rpl13
Rpl19
Rplp0
Rplp1
Chchd2
Chchd10
Cox8a
Atp1a1
Rpl32
Rps20
Rpl18a
Rpl39
Rpl37
Rpl7
Rps8
Rps23
...
2
1
0
-1
-2
Malat1
Neat1
Son
Gls
Zbtb20
Vegfa

G

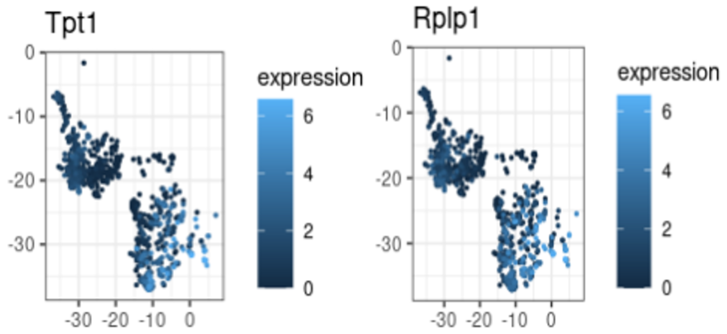
Tpt1
expression
6
4
2
0
0
-10
-20
-30
-30 -20 -10 0
Rplp1
expression
6
4
2
0
0
-10
-20
-30
-30 -20 -10 0

Figure S1

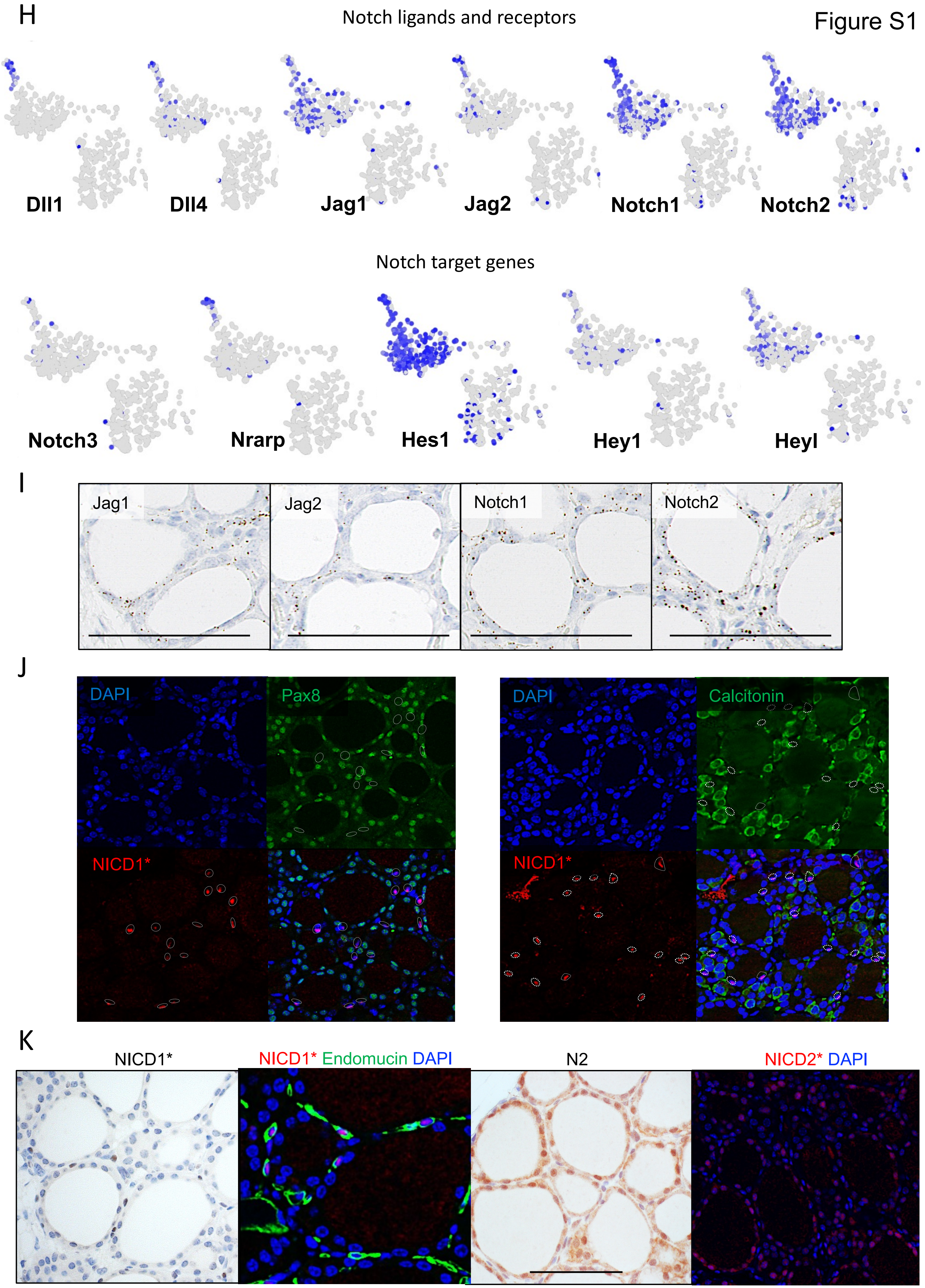
H
Notch ligands and receptors
Dll1
Dll4
Jag1
Jag2
Notch1
Notch2
Notch target genes
Notch3
Nrarp
Hes1
Hey1
Heyl
I
Jag1
Jag2
Notch1
Notch2
J
DAPI
Pax8
NICD1*
DAPI
Calcitonin
NICD1*
K
NICD1*
NICD1* Endomucin DAPI
N2
NICD2* DAPI

#### Figure S2. Notch inhibition induces thyrocyte defects

**(A)** Representative immunofluorescent detection of NICD1*, plus quantification of NICD1* and NICD2*, in thyroid sections from mice (n = 5-6) treated as in Fig. 2A. 40X objective. Ave. ± s.d.

**(B)** Quantification of HeyL, Hey1 and Nrarp mRNA expression detected using RNAscope probes in thyroid sections from mice (n = 4) treated as in Fig. 2B. Percentage of positive cells and H score, calculated by totaling the percentage of cells in each bin, according to the number of dots/cell (0, 0 dots; 1, 1-3 dots; 2, 4-9 dots; 3, 10-15 dots and 4, >15 dots). Ave. ± s.d.

**(C)** Notch3 mRNA expression detected using RNAscope in thyroid sections from mice (n = 3) treated as in (B), showing sporadic Notch3 expression primarily in non-follicular cells. Scale bars = 100mm. Ave. H score ± s.d.

**(D)** NES of a Notch target gene set (Hey1, Nrarp, HeyL, Notch3) from thyroids isolated at the indicated times after treating mice (n = 5) as in (A).

**(E)** Top 30 pathways downregulated following aJ12 treatment (relative to isotype control, treated as in A, n = 5 mice) obtained using GSEA.

**(F)** RNA sequencing (nRPKM fold change) of the indicated thyrocyte and parafollicular (PFC) marker genes from thyroids isolated from mice (n = 5) treated as in (A). Ave. ± s.d.

**(G)** qRT-PCR using Taqman probes (fold change in expression normalized to actin mRNA) of the indicated marker genes from thyroids isolated from mice (n = 6) treated as in (A) for 5 days. Ave ± s.d.

**(H)** Representative immunohistochemical detection of calcitonin and calcitonin gene-related protein (CGRP) in thyroid sections from mice (n = 5) treated as in (A) for 5 days. Scale bars = 100mm.

**(I)** Percentage of Pax8+ cells in the thyroids of mice (n = 4) treated as in (A) and assessed as in Sup. Fig. S1E. Ave. ± s.d.

**(J)** Distribution of expression of the indicated genes in TFC2 and TFC3 cells from mice (n = 3) treated as in (B).

**(K)** Representative immunofluorescent detection (from n = 4 wells) of Pax8 in FRTL5 cells treated for 2 days with aRW (25µg/ml), aJ12 (25µg/ml each) or aN123 (25µg/ml each). aJ12 decreases the number of cells with nuclear Pax8 and the overall intensity of Pax8 staining whereas aN123 nearly completely eliminates nuclear Pax8 staining. DAPI was used for nuclear staining. 20X objective.

**(L)** Representative immunofluorescent images of the thyroid organoids isolated from mice treated as in (A) and stained for Nkx2-1 (red), calcitonin (green) and Pax8 (white). 20X objective.

**(M)** Number of organoids generated from 10000 single cells isolated from thyroids of mice treated as in (A). Single cells were either isolated from 7 thyroids, digested in a pool, and plated into 3 wells, before counting organoids/well on day 6 (D6) or isolated from 8 thyroids, pooled into 2 thyroids/pool, and plated into single wells before counting organoids/well on day 7 (D7). Ave. ± s.d.; n = 3-4 wells.

**(N)** aJ12 alters multiple characteristics of thyroid organoids. Jag blockade reduced the percentage of organoids that were all positive for Pax8 immunofluorescence while increasing the percentages of those that were all negative or mixed (Pax8+ and Pax8- cells in an organoid). Jag blockade also led to reductions in organoid size (small<20 cells/organoid≤big) and development (developed, > 1 follicle formed; underdeveloped, no follicles, unorganized cell mass). n = 49 and 33 organoids from mice treated with aRW or aJ12, respectively.

**(O)** GSEA of Reactome pathways from cells in TFC3, highlighting "posttranslational protein modification" with NES < -2 in both the aN12/aRW and aJ12/aRW comparisons.

**(P)** Akaike Information Criterion (AIC) and Bayesian Information Criterion (BIC) statistics per number of topics represented in the single cell data.

**(Q)** tSNE plots of the thyrocyte clusters showing a blue heatmap of each topic, plus boxplots showing the topic scores in each treatment group. Notch blockade most notably affected Topic 1, decreasing the score, and Topic 2, increasing the score. n=3 thyroids/group.

Statistical significance was assessed using the unpaired two-tailed Student's t-test with Welch's correction: $p<0.05$, *; $p<0.01$, **; $p<0.001$, ***; $p<0.0001$, ****. See also Figure 2.

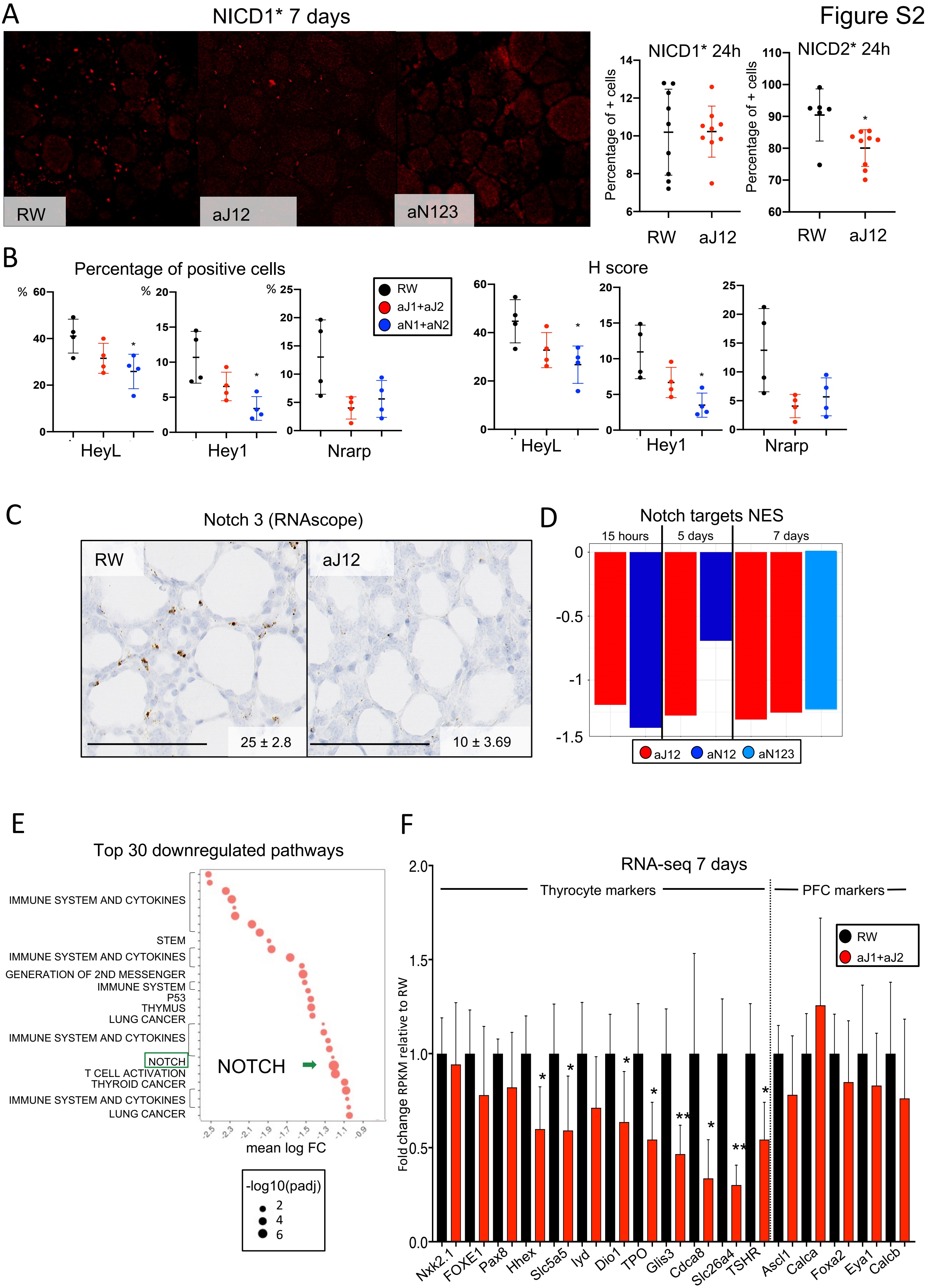
Figure S2
A
NICD1* 7 days
RW
aJ12
aN123
NICD1* 24h
NICD2* 24h
Percentage of + cells
RW
aJ12
B
Percentage of positive cells
H score
RW
aJ1+aJ2
aN1+aN2
HeyL
Hey1
Nrarp
C
Notch 3 (RNAscope)
RW
aJ12
25 ± 2.8
10 ± 3.69
D
Notch targets NES
15 hours
5 days
7 days
aJ12
aN12
aN123
E
Top 30 downregulated pathways
IMMUNE SYSTEM AND CYTOKINES
STEM
IMMUNE SYSTEM AND CYTOKINES
GENERATION OF 2ND MESSENGER
IMMUNE SYSTEM
P53
THYMUS
LUNG CANCER
IMMUNE SYSTEM AND CYTOKINES
NOTCH
T CELL ACTIVATION
THYROID CANCER
IMMUNE SYSTEM AND CYTOKINES
LUNG CANCER
NOTCH
mean log FC
-log10(padj)
F
RNA-seq 7 days
Thyrocyte markers
PFC markers
RW
aJ1+aJ2
Fold change RPKM relative to RW
Nkx2.1
FOXE1
Pax8
Hhex
Slc5a5
Iyd
Dio1
TPO
Glis3
Cdca8
Slc26a4
TSHR
Ascl1
Calca
Foxa2
Eya1
Calcb

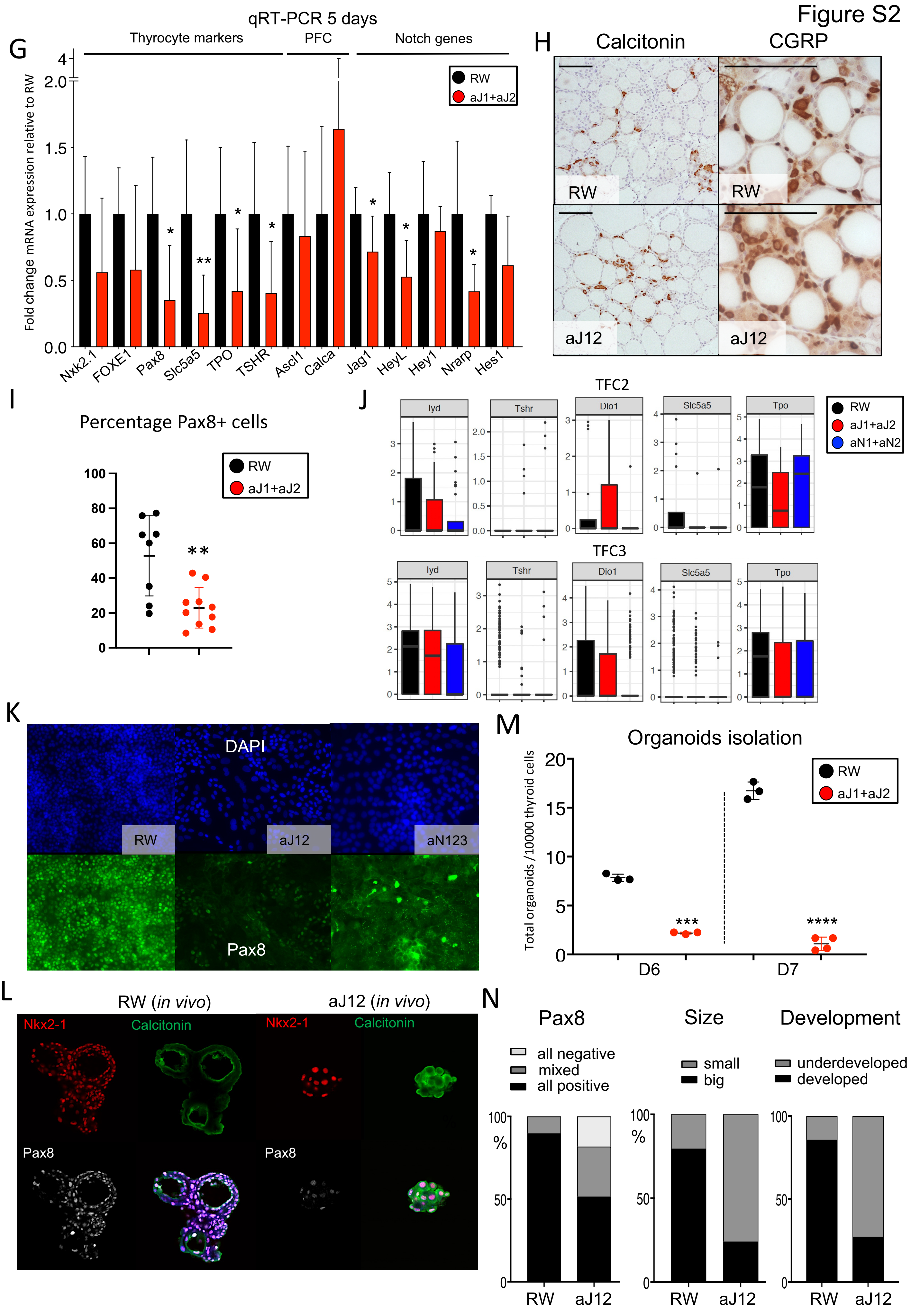
Figure S2
G
qRT-PCR 5 days
Thyrocyte markers
PFC
Notch genes
Fold change mRNA expression relative to RW
RW
aJ1+aJ2
Nkx2.1
FOXE1
Pax8
Slc5a5
TPO
TSHR
Ascl1
Calca
Jag1
HeyL
Hey1
Nrarp
Hes1
H
Calcitonin
CGRP
RW
aJ12
I
Percentage Pax8+ cells
RW
aJ1+aJ2
J
TFC2
TFC3
Iyd
Tshr
Dio1
Slc5a5
Tpo
RW
aJ1+aJ2
aN1+aN2
K
DAPI
RW
aJ12
aN123
Pax8
M
Organoids isolation
Total organoids /10000 thyroid cells
RW
aJ1+aJ2
D6
D7
L
RW (in vivo)
aJ12 (in vivo)
Nkx2-1
Calcitonin
Pax8
N
Pax8
Size
Development
all negative
mixed
all positive
small
big
underdeveloped
developed
%
RW
aJ12

### Figure S2

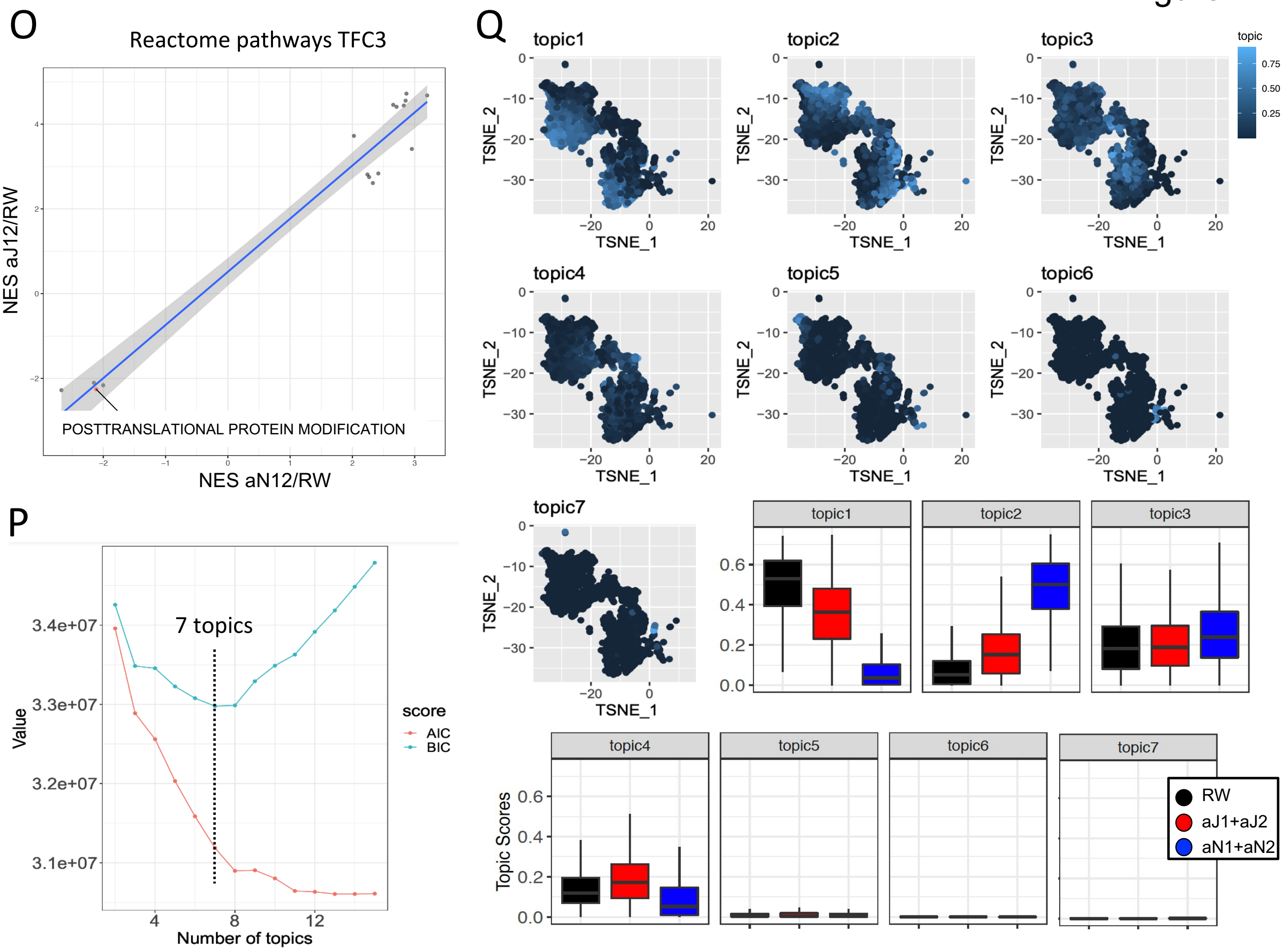
O
Reactome pathways TFC3
NES aJ12/RW
NES aN12/RW
POSTTRANSLATIONAL PROTEIN MODIFICATION
P
7 topics
Value
Number of topics
score
AIC
BIC
Q
topic1
topic2
topic3
topic4
topic5
topic6
topic7
TSNE_1
TSNE_2
Topic Scores
RW
aJ1+aJ2
aN1+aN2

**Figure S3. Notch blockade induces mitochondrial defects in thyrocytes**

(A) Distribution of expression of the indicated mitochondrial genes in TFC1 cells isolated from mice (n = 3) treated as in Fig. 3A.
(B) Representative scanning electron microscopy images of mitochondria from non-thyrocyte cells from thyroids of mice (n = 3) treated as in (A). Scale bars = 100nm.
(C) Fluorescent intensity of TMRM in primary thyrocytes from mice (n = 5) treated as in (A).
(D) Quantification of Cox staining using the VitroView$^{TM}$ kit in thyroid cryosections of mice (n =4) treated as in (A). Ave. ± s.d.
(E) Cytochrome c oxidation kinetics in primary thyrocytes isolated from mice (n = 5) treated as in (A). Ave. ± s.d.
(F) Ratio of mitochondrial DNA (mDNA, Nd1 and 16S) versus genomic DNA (gDNA, HK and 18S) in thyroids from mice (n = 9) treated as in (A). Ave. ± s.d.
(G) Seahorse assay of ATP production rates in the FRTL5 thyrocyte cell line cultured for 2 days as in Fig. 3E. Ave ± s.d.; n = 13 wells.
(H) Percentages of thyrocytes with high (≥ 2500 MFI) or low mitotracker (<2500 MFI) intensities or mitosox positivity (≥ 450 MFI) as in Fig. 3F. Ave ± s.d.; n = 5 mice.
(I) Mean fluorescence intensities of mitotracker and mitosox in cells from (G) with low mitotracker staining. Ave. ± s.d.; n = 5 mice.
(J) Mean fluorescence intensity following DCFDA staining to measure ROS levels as in Fig. 3H.
(K) Principal component analysis (PCA) of thyroid metabolites from mice (n = 4) treated as in (A).
(L) Quantification of glycolysis and TCA cycle metabolites of samples in (J). Succinate, Citrate, Lactate and Pyruvate were quantified using targeted analysis (ng/g), and glucose-6-phosphate levels are reported as MS peak area units. Ave. ± s.d.; n = 4 thyroids.
(M) Quantification of ketogenesis metabolites of samples in (J). 3-OH-butyrate and 2-OH-3-methylbutyric acid were quantified using targeted analysis or with MS peak area units, respectively. Ave ± s.d.; n = 4 thyroids.
(N) Quantification in MS peak area units of urea cycle and glutamine cycle metabolites of samples in (J). Ave. ± s.d.; n = 4 thyroids.
(O) Quantification of catalase activity in thyrocytes isolated from mice (n = 5) treated as in (A). Ave. ± s.d.
(P) Quantification of glutathione disulfide (GSSG) in thyrocytes isolated from mice (n = 5) treated as in (A). Ave. ± s.d.
(Q) T4 levels in the media of FRTL5 thyrocyte cell cultures as in Fig. 3E. Ave. ± s.d.; n = 6 wells.

Statistical significance was assessed using the unpaired two-tailed Student's t-test with Welch's correction: $p<0.05$, *; $p<0.01$, **; $p<0.001$, ***; $p<0.0001$, ****. See also Figure 3.

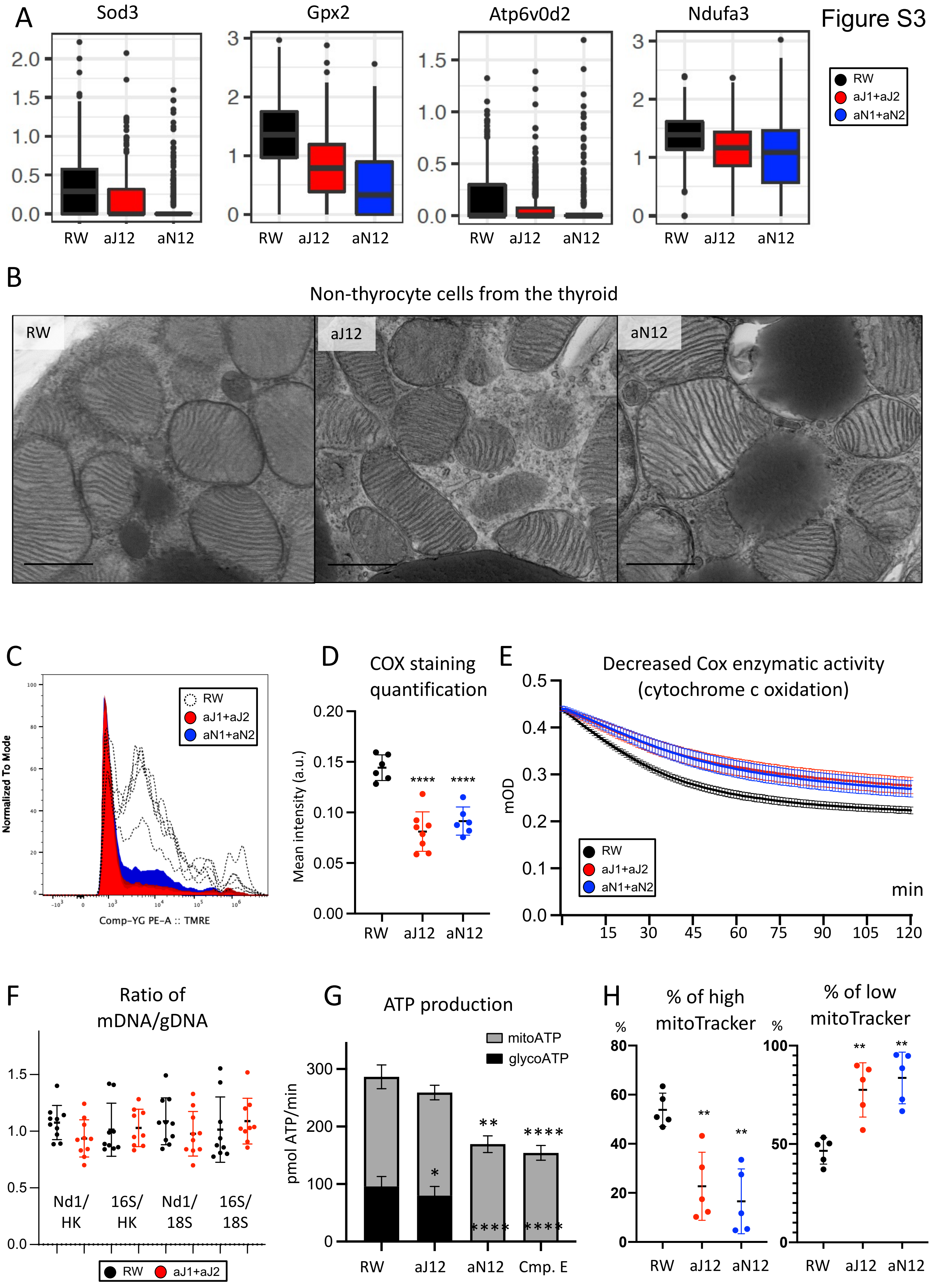

A
Sod3
Gpx2
Atp6v0d2
Ndufa3
RW
aJ1+aJ2
aN1+aN2
RW aJ12 aN12
B
Non-thyrocyte cells from the thyroid
RW
aJ12
aN12
C
Normalized To Mode
Comp-YG PE-A :: TMRE
D
COX staining quantification
Mean intensity (a.u.)
****
E
Decreased Cox enzymatic activity (cytochrome c oxidation)
mOD
min
F
Ratio of mDNA/gDNA
Nd1/ HK
16S/ HK
Nd1/ 18S
16S/ 18S
G
ATP production
mitoATP
glycoATP
pmol ATP/min
RW aJ12 aN12 Cmp. E
H
% of high mitoTracker
% of low mitoTracker
%


Figure S3

Figure S3

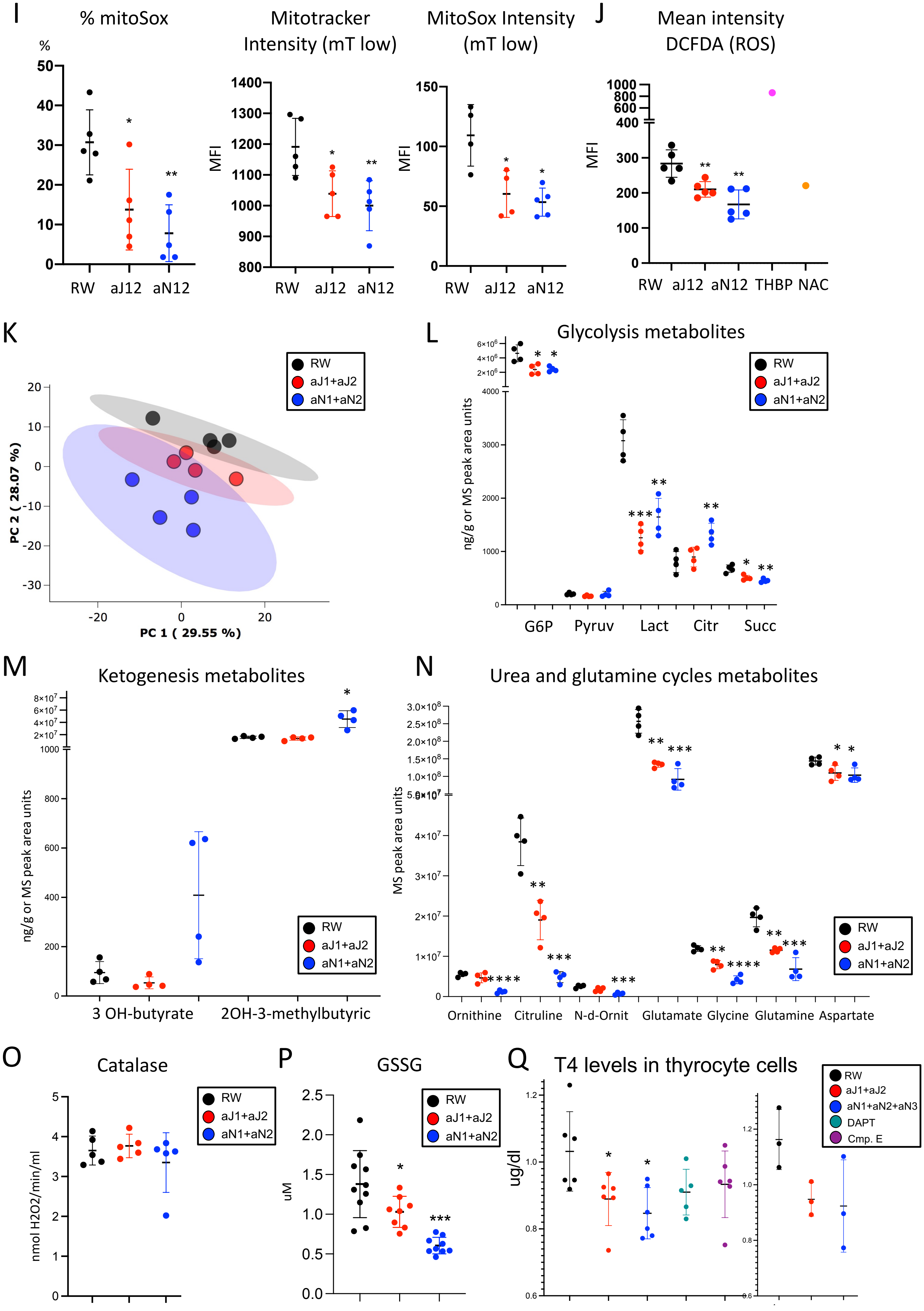

I
% mitoSox
%
RW aJ12 aN12
Mitotracker Intensity (mT low)
MFI
MitoSox Intensity (mT low)
MFI
J
Mean intensity DCFDA (ROS)
MFI
RW aJ12 aN12 THBP NAC
K
PC 2 ( 28.07 %)
PC 1 ( 29.55 %)
RW
aJ1+aJ2
aN1+aN2
L
Glycolysis metabolites
ng/g or MS peak area units
G6P Pyruv Lact Citr Succ
RW
aJ1+aJ2
aN1+aN2
M
Ketogenesis metabolites
ng/g or MS peak area units
3 OH-butyrate 2OH-3-methylbutyric
RW
aJ1+aJ2
aN1+aN2
N
Urea and glutamine cycles metabolites
MS peak area units
Ornithine Citruline N-d-Ornit Glutamate Glycine Glutamine Aspartate
RW
aJ1+aJ2
aN1+aN2
O
Catalase
nmol H2O2/min/ml
RW
aJ1+aJ2
aN1+aN2
P
GSSG
uM
RW
aJ1+aJ2
aN1+aN2
Q
T4 levels in thyrocyte cells
ug/dl
RW
aJ1+aJ2
aN1+aN2+aN3
DAPT
Cmp. E

**Figure S4. Notch inhibition triggers hypothyroidism and a cascade of thermoregulation defects dependent on thyroid hormones**

(A) Quantification of thyroid histological changes assessed by percentage of thyroid samples, n=10 thyroids per group. Follicles with attenuated epithelium and less than 4 colloid resorption vacuoles were considered inactive. Thyroid sections estimated to have more than 40% of peripheral follicles inactive were interpreted to have reduced activity. Thyroid sections with all follicles lined by cuboidal epithelium were scored as 'none' (white); thyroid with <10% follicles lined by flat epithelium were scored as 'minimal' (light grey); 10-40% follicles = 'mild' (grey); 40-75% follicles = 'moderate' (dark grey) and thyroids with >80% follicles lined by flat epithelium were scored as 'marked' (black). Mice were treated as in Fig 4A.

(B) Serum levels of triiodothyronine (T3) and thyroxine (T4) measured at 7 days post treatment. Mice were treated with a single i.p. dose of aRW (black; 40 mg/kg); aJ12 (red; 20 mg/kg each); aN123 (blue, 5, 10 and 20 mg/kg); aN1 (yellow, 5 mg/kg); aN2 (green, 10 mg/kg); aN3 (brown, 20 mg/kg); aJ1 (turquoise, 20 mg/kg) or aJ2 (mustard, 20 mg/kg); aN12 (dark blue; 5,10 mg/kg), aN23 (pink; 10, 20 mg/kg, aN3 twice a week) and aN13 (purple; 5, 20 mg/kg, aN3 injected twice a week). Ave. ± s.d; n $\geq$ 5 mice.

(C) Serum levels of Thyrotropin Releasing Hormone (TRH), secreted by the hypothalamus, in mice treated as in (A) for 5 days. Ave. ± s.d.; n = 6 mice.

(D) Serum levels of Thyrotropin Secreting Hormone (TSH), secreted by the pituitary gland, in mice treated as in (A) for 5 or 7 days. Ave. ± s.d.; n = 6 mice.

(E) Serum levels of calcitonin in mice treated as in (A) for 7 days. Ave. ± s.d.; n $\geq$ 4 mice.

(F) T4 serum levels assessed by ELISA in Ob/Ob or WT mice housed at the indicated temperatures and treated for 7 days as in (A). Ave. ± s.d.; n $\geq$ 4 mice.

(G) T4 levels following T3 and T4 administration (1 µg/g and 0.05 µg/g, respectively) according to the schematic. Ave. ± s.d.; n = 4 mice.

(H) Body temperatures measured using a rectal probe in mice treated as in (G) and housed at 22°C. Ave. ± s.d.; n = 5 mice.

(I) Body temperatures measured using a rectal probe in mice treated as in (G) and housed at 4°C. Ave ± s.d.; n = 5 mice.

(J) Electron microscopy images of mitochondria of thyrocytes from mice treated as in (G) for 7 days, n = 3 mice. Arrows = aberrant mitochondria. Scale bars = 400nm.

(K) Representation of the temperature dissipated from the tail measured by infrared thermography in mice treated as in (G) during 6 days. Ave. ± s.d.; n = 5 mice.

(L) Score of the BAT activity, assessed blindly in the BAT sections from mice treated as in (G). Score of 1 is white adipose tissue phenotype and 10 BAT at its maximum activity. Ave. ± s.d.; n=5 mice.

(M) Picture of the interscapular brown adipose tissue (BAT) of mice treated as in (A) for 7 days, n = 5 mice.

(N) Fold RPKM of the indicated genes in the BAT of mice treated as in (G). Ave. ± s.d.; n = 5 mice.

(O) Immunohistochemistry against UCP1 in BAT from mice treated for 5 days as in (A), n = 5 mice. Scale bars = 100mm.

(P) BAT temperature in mice treated as in (G). Temperature was measured by IPT transponders implanted in the interscapular region. Data were collected from 3 independent experiments. Ave. ± s.d.; n $\geq$ 10 mice.

(Q) Serum levels of glucose, cholesterol and triglycerides of mice treated as in (G). Ave. ± s.d.; n = 5 mice.
(R) NEFA serum levels of mice treated as in (G). Ave. ± s.d.; n = 5 mice.

Statistical significance was assessed using the unpaired two-tailed Student's t-test with Welch's correction: $p<0.05$, *; $p<0.01$, **; $p<0.001$, ***; $p<0.0001$, ****. See also Figure 4.

#### Figure S4

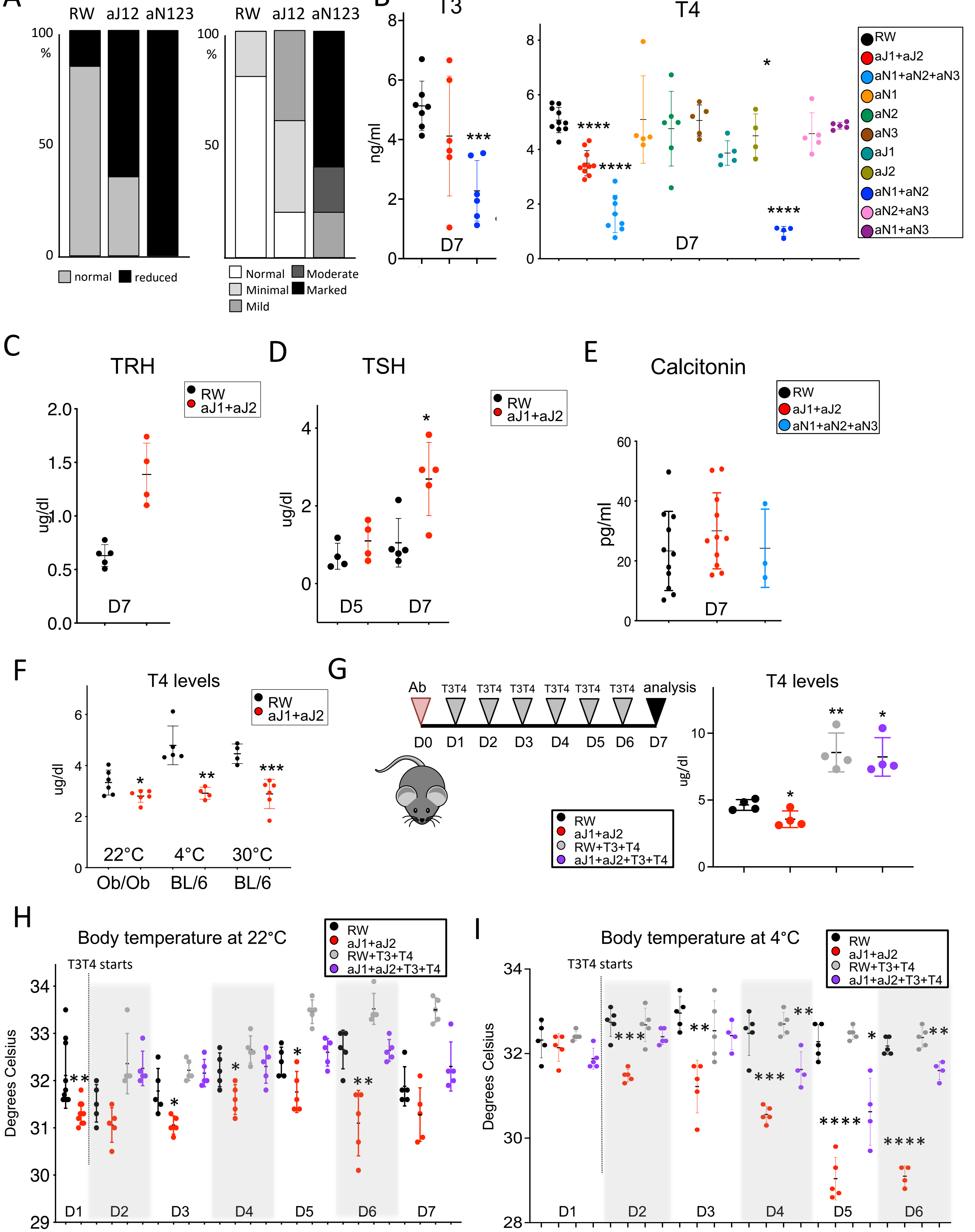
A
Thyroid activity
RW aJ12 aN123
100 %
50
0
normal reduced
Flattening
RW aJ12 aN123
Normal Moderate
Minimal Marked
Mild
B
T3
ng/ml
D7
***
T4
D7
****
*
RW
aJ1+aJ2
aN1+aN2+aN3
aN1
aN2
aN3
aJ1
aJ2
aN1+aN2
aN2+aN3
aN1+aN3
C
TRH
ug/dl
D7
RW
aJ1+aJ2
D
TSH
D5
D7
*
E
Calcitonin
pg/ml
D7
F
T4 levels
ug/dl
22°C 4°C 30°C
Ob/Ob BL/6 BL/6
G
Ab T3T4 T3T4 T3T4 T3T4 T3T4 T3T4 analysis
D0 D1 D2 D3 D4 D5 D6 D7
RW
aJ1+aJ2
RW+T3+T4
aJ1+aJ2+T3+T4
T4 levels
ug/dl
H
Body temperature at 22°C
T3T4 starts
Degrees Celsius
D1 D2 D3 D4 D5 D6 D7
I
Body temperature at 4°C
T3T4 starts
Degrees Celsius
D1 D2 D3 D4 D5 D6

Figure S4

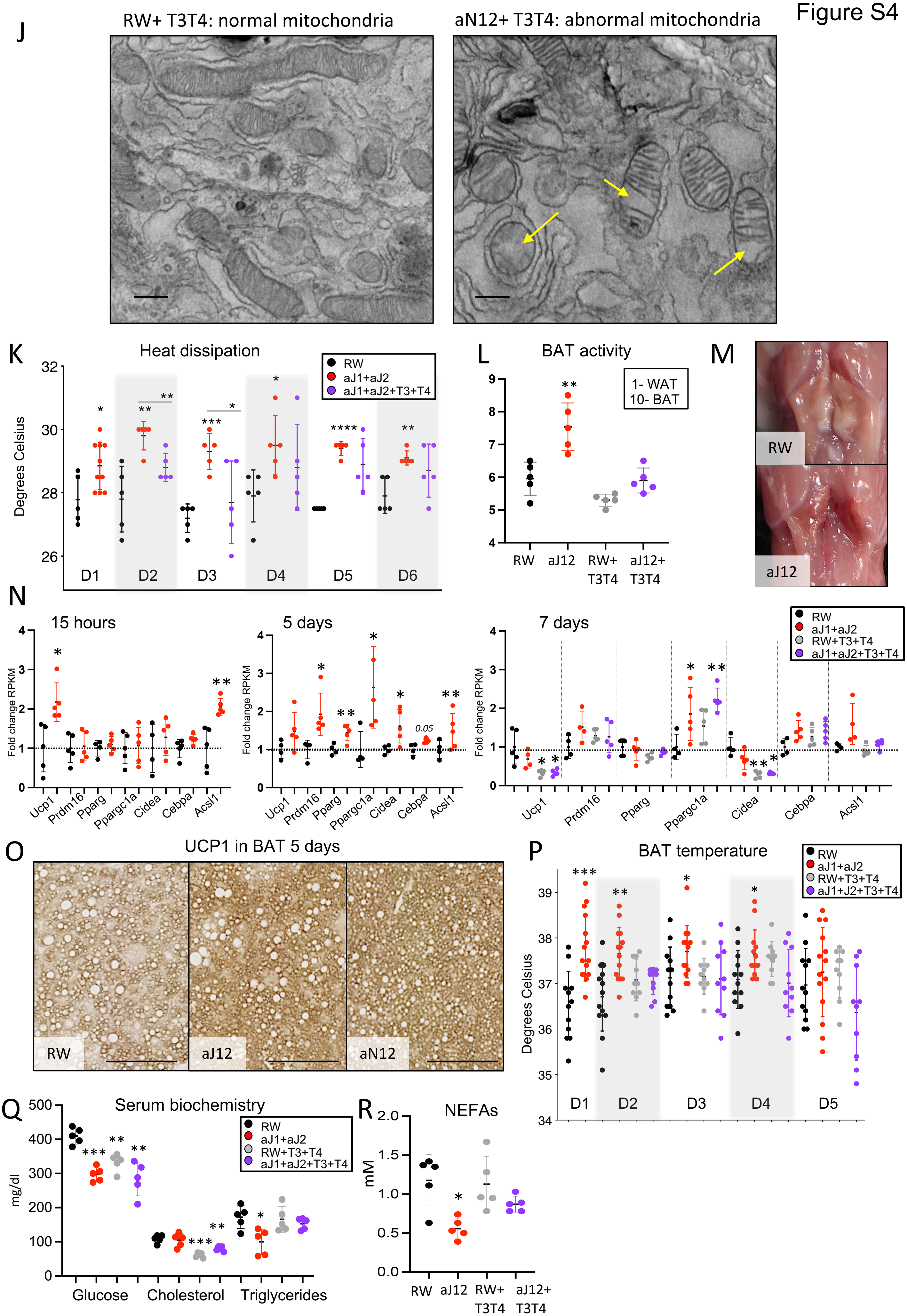
J
RW+ T3T4: normal mitochondria
aN12+ T3T4: abnormal mitochondria
K
Heat dissipation
Degrees Celsius
RW
aJ1+aJ2
aJ1+aJ2+T3+T4
D1
D2
D3
D4
D5
D6
L
BAT activity
1- WAT
10- BAT
RW
aJ12
RW+ T3T4
aJ12+ T3T4
M
RW
aJ12
N
15 hours
5 days
7 days
Fold change RPKM
0.05
RW
aJ1+aJ2
RW+T3+T4
aJ1+aJ2+T3+T4
Ucp1
Prdm16
Pparg
Ppargc1a
Cidea
Cebpa
Acsl1
O
UCP1 in BAT 5 days
RW
aJ12
aN12
P
BAT temperature
Degrees Celsius
RW
aJ1+aJ2
RW+T3+T4
aJ1+J2+T3+T4
D1
D2
D3
D4
D5
Q
Serum biochemistry
mg/dl
RW
aJ1+aJ2
RW+T3+T4
aJ1+aJ2+T3+T4
Glucose
Cholesterol
Triglycerides
R
NEFAs
mM
RW
aJ12
RW+ T3T4
aJ12+ T3T4

#### Figure S5. Notch genetic alterations induce hypothyroidism in mice and patients

(A) Scheme of tamoxifen administration in $N2^{fl/fl}$; Tg-$CreER^{T2}$+/+ mice and their respective Tg-$CreER^{T2}$-/- littermates. After two cycles of tamoxifen, mice were administered 5 mg/kg of RW or aN1 antibody and tissues were collected 7 days after antibody treatment.
(B) Gel showing PCR products of gDNA isolated from thyroids of mice from (A) and amplified using primers to detect the WT Notch2 band (1485bp) or the recombined allele (480bp).
(C) Immunohistochemistry against Cre (up) or Notch2 (down) on thyroid sections from mice from (A), n = 5 mice. Scale bars = 100mm.
(D) Fold expression of the represented genes analyzed by qRT-PCR using Taqman probes in the thyroid of mice treated as in (A). Gene expression was normalized to actin and is shown as relative to $N2^{Fl/Fl}$ neg+aRW. Ave. ± s.d.; n = 5 mice.
(E) Infrared thermography pictures of mice treated as in (A) and measured at 4 days after treatment. The bar on the right shows the maximum and minimum temperature registered in each picture with the corresponding gradient of colors, n = 5 mice.
(F) Picture of the interscapular brown adipose tissue (BAT) of mice treated as in (A) for 7 days, n= 5 mice.
(G) Serum levels of glucose, cholesterol and triglycerides of mice treated as in (A). Ave. ± s.d.; n$\geq$ 4 mice.
(H) Double immunofluorescence against cleaved (active) NICD1* and endomucin in thyroids from (A), n = 5 mice. Images were taken using 40X objective.

Statistical significance was assessed using the unpaired two-tailed Student's t-test with Welch's correction: p<0.05, *; p<0.01, **; p<0.001, ***; p<0.0001, ****. See also Figure 5.

### Figure S5

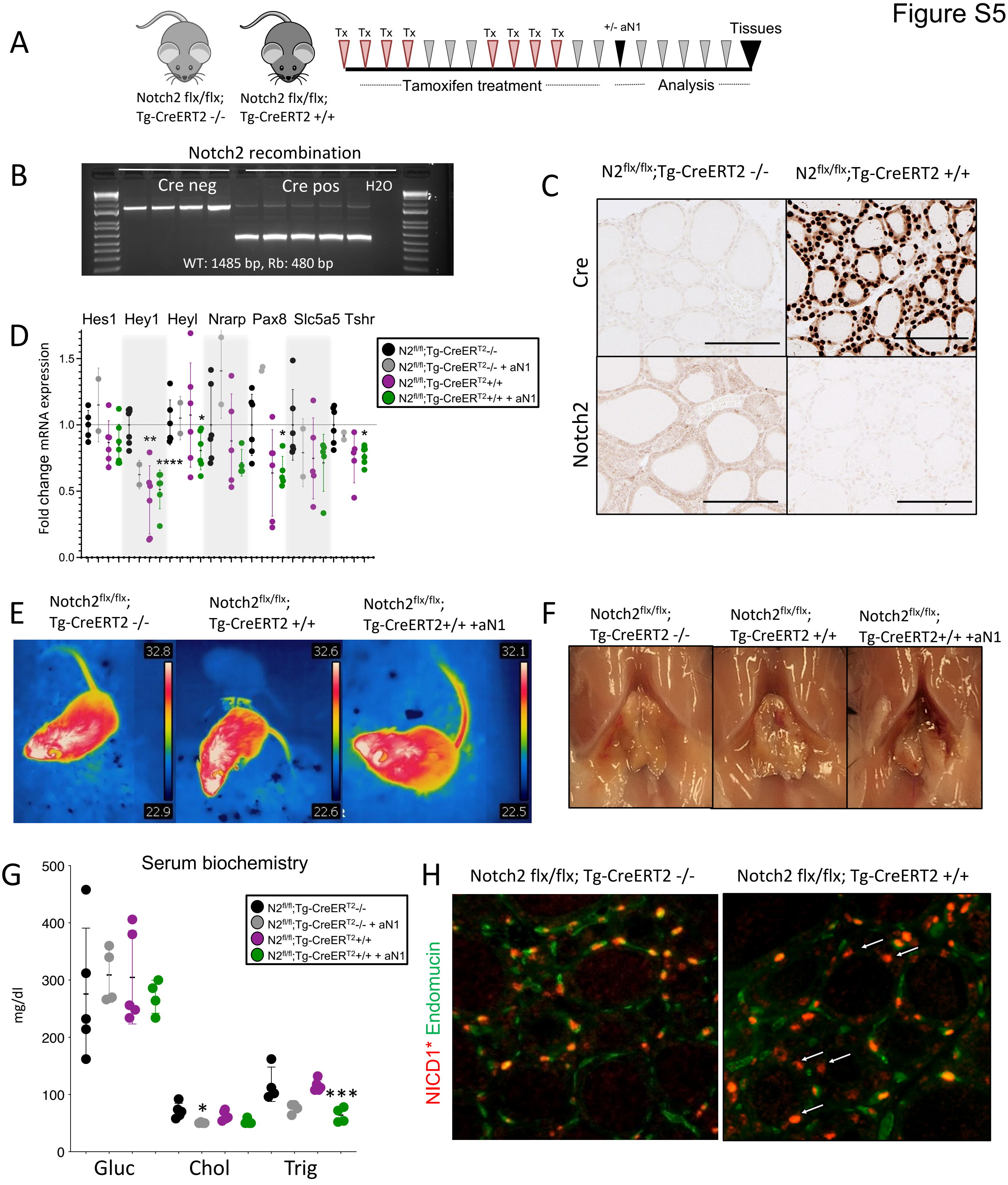
A
Notch2 flx/flx; Tg-CreERT2 -/-
Notch2 flx/flx; Tg-CreERT2 +/+
Tx Tx Tx Tx Tx Tx Tx Tx
+/- aN1
Tissues
Tamoxifen treatment
Analysis
B
Notch2 recombination
Cre neg
Cre pos
H2O
WT: 1485 bp, Rb: 480 bp
C
N2flx/flx;Tg-CreERT2 -/-
N2flx/flx;Tg-CreERT2 +/+
Cre
Notch2
D
Hes1 Hey1 Heyl Nrarp Pax8 Slc5a5 Tshr
Fold change mRNA expression
N2fl/fl;Tg-CreERT2-/-
N2fl/fl;Tg-CreERT2-/- + aN1
N2fl/fl;Tg-CreERT2+/+
N2fl/fl;Tg-CreERT2+/+ + aN1
E
Notch2flx/flx; Tg-CreERT2 -/-
Notch2flx/flx; Tg-CreERT2 +/+
Notch2flx/flx; Tg-CreERT2+/+ +aN1
32.8 22.9 32.6 22.6 32.1 22.5
F
Notch2flx/flx; Tg-CreERT2 -/-
Notch2flx/flx; Tg-CreERT2 +/+
Notch2flx/flx; Tg-CreERT2+/+ +aN1
G
Serum biochemistry
mg/dl
Gluc
Chol
Trig
N2fl/fl;Tg-CreERT2-/-
N2fl/fl;Tg-CreERT2-/- + aN1
N2fl/fl;Tg-CreERT2+/+
N2fl/fl;Tg-CreERT2+/+ + aN1
H
Notch2 flx/flx; Tg-CreERT2 -/-
Notch2 flx/flx; Tg-CreERT2 +/+
NICD1* Endomucin
